# Tumor-Associated Macrophages Promote Brain Metastasis

**DOI:** 10.64898/2026.06.29.735286

**Authors:** Chaitali Khan, Nasser M Rusan

## Abstract

Brain metastasis affects 20-40% of cancer patients and remains largely incurable, yet how metastatic cells engage and remodel brain-barrier interfaces is poorly understood. This gap stems in large part from the scarcity of genetically tractable *in vivo* systems that can resolve tumor-host interactions in their native context. Although tumor-associated macrophages (TAMs) dominate the brain metastatic microenvironment, their causal contribution to colonization has been difficult to establish. Here, we develop the adult *Drosophila* brain as a platform to address these questions, using allograft transplantation of *lgl*^-/-^ neural stem cell-derived tumors. We find that tumors colonize the brain surface and deform it through collective, sheet-like invasion, recapitulating features of human leptomeningeal disease without parenchymal infiltration. While tumors compromise both functional and structural integrity of blood-brain barrier, they fail to breach the basement membrane (BM), which we identify as the principal barrier restricting invasion. Critically, tumors recruit TAMs to the metastasized tumor, and genetic depletion of TAMs markedly reduces brain metastasis without affecting initial dissemination. This establishes a causal role for macrophages in brain colonization, potentially through BM remodeling. Together, these findings reveal a conserved, macrophage-dependent mechanism of brain metastasis and provide a genetically accessible model that fills a critical gap in the field.

## INTRODUCTION

Dissecting the complex interactions within the tumor microenvironment (TME), which encompasses tumor cells, infiltrating stromal, immune populations, and host organs, remains a major challenge for achieving a mechanistic understanding of tumor metastasis (Grant & Ferrer, 2025; Joyce & Pollard, 2009; Quail & Joyce, 2013). A key limitation in the field is the lack of genetically tractable *in vivo* systems that allow mechanistic interrogation of tumor-host interactions at cellular resolution within a native tissue context. *Drosophila melanogaster* has long contributed to the understanding of conserved genes and signaling pathways underlying tumor growth and metastasis (Choutka et al., 2022; Khan & Rusan, 2024; Mirzoyan et al., 2019; Rudrapatna et al., 2012; Villegas et al., 2019), and has more recently emerged as a powerful *in vivo* platform for studying TME interactions, particularly the role of innate immunity in tumor progression (Bangi, 2013; Bilder et al., 2021; Khan & Rusan, 2024; Sharpe et al., 2023; Teles-Reis & Rusten, 2026; Wang et al., 2014). Critically, the absence of adaptive immunity in *Drosophila* provides a clean genetic context in which to isolate the contribution of innate immune cells to tumor progression. Our recent work extended this framework to the adult stage by establishing a reproducible allograft-based model, enabling the study of tumor-host interactions during metastatic colonization of distant organs (Khan & Rusan, 2025).

Brain metastasis is the most commonly diagnosed intracranial malignancy, occurring in 20-40% of all cancer patients, with metastatic colonization arising in two major anatomically distinct compartments (Kamp et al., 2018; Lamba et al., 2021; Shojania et al., 2003). The first is Parenchymal Metastasis, the most common form, requires tumor cells to invade the neural tissue by breaching the blood-brain barrier (BBB), a multi-component interface comprising tight junction-sealed endothelial cells, pericytes, astrocytic endfeet, and an underlying basement membrane (BM) (Arvanitis et al., 2020; Steeg, 2021). The second is Leptomeningeal Disease (LMD), which involves spreading of tumor cells to the cerebrospinal fluid (CSF) in the subarachnoid space and colonization of the pia mater, requiring tumor cells to traverse the arachnoid-CSF barrier that is formed by tight junction-connected arachnoid fibroblast-like cells (Freret & Boire, 2024; Ozair et al., 2025). Regardless of type, a central and unresolved question across all forms of brain metastasis is how tumor cells interact with, and remodel, the brain-barrier interfaces during metastatic invasion.

The *Drosophila* brain, although lacking a closed circulatory vasculature, is encased by a multi-layered barrier system that is anatomically analogous to the mammalian meningeal layers and functionally analogous to both the mammalian BBB and arachnoid-CSF barrier (Contreras & Klambt, 2023; Limmer et al., 2014; Love & Dauwalder, 2019). The outermost layer, the neural lamella (NL) or BM, is an acellular extracellular matrix (ECM) sheath in direct contact with the circulating hemolymph. Beneath the NL, the perineurial glia (PNG) form the outermost cellular layer (Kremer et al., 2017; Pogodalla et al., 2022). The subperineurial glia (SPG), lying beneath the PNG, form the primary paracellular barrier through pleated septate junctions (SJs), the functional equivalent of vertebrate tight junctions. Thus, the SJs completely seal the nervous system from the hemolymph and block the paracellular entry of macromolecules and cells (Bainton et al., 2005; Kremer et al., 2017; Stork et al., 2008; Tepass & Hartenstein, 1994). Our recent demonstration that metastatic *lgl*^-/-^ NSC-derived tumors form a close association with the outer barrier layers of the adult *Drosophila* brain (Khan & Rusan, 2025) proves to be a tractable *in vivo* model for dissecting the cellular mechanisms by which metastatic tumor cells interact with, and remodel brain-barrier interfaces.

Among the diverse cellular players shaping tumor-host interactions, tumor-associated macrophages (TAMs) emerge as key regulators of cancer progression from tumor initiation to metastatic colonization at distant organ sites, exhibiting a range of functional roles shaped by their plasticity and local microenvironmental cues (Friedman-DeLuca et al., 2024; Mantovani et al., 2024; Murray et al., 2014; Pittet et al., 2022; Qian & Pollard, 2010). At the primary site, TAMs promote metastatic dissemination by inducing epithelial-to-mesenchymal transition (EMT) (Bonde et al., 2012; Li et al., 2022; Su et al., 2014), remodeling the ECM (Goswami et al., 2005; Wyckoff et al., 2004), and facilitating intravasation at tumor microenvironment of metastasis (TMEM) doorways (Condeelis & Pollard, 2006; Entenberg et al., 2018; Harney et al., 2015; Robinson et al., 2009). Similarly, at distant metastatic sites, TAMs prime the pre-metastatic niche (Kitamura et al., 2015; Peinado et al., 2017; Sharma et al., 2015), aid extravasation of circulating tumor cells by increasing endothelial permeability (Qian et al., 2009; Qian et al., 2011), and support early colonization through ECM remodeling and suppression of tumoricidal immunity (Doedens et al., 2010; Hiratsuka et al., 2002; Qian et al., 2009).

The brain metastasis TME is dominated by TAMs, comprising two ontogenetically distinct populations: brain-resident microglia and recruited monocyte-derived macrophages (MDMs), as well as a less well-characterized population of border-associated macrophages (BAMs) residing at perivascular and meningeal interfaces (Friebel et al., 2020; Gonzalez et al., 2022; Karimi et al., 2023; Klemm et al., 2020; Ratzabi et al., 2026). Microglia predominate during the early stages at the tumor-brain interface and contribute to extravasation through remodeling of the BBB (Gan et al., 2024; Lorger & Felding-Habermann, 2010; Pukrop et al., 2010). In contrast, MDMs are progressively recruited from the circulation, accumulate within the tumor core, less so at the brain-tumor interface, and are known to promote tumor expansion by modulating immune response and remodeling barrier structure and ECM (Friebel et al., 2020; Gan et al., 2024; Klemm et al., 2020). Despite their abundance and functional prominence, the cellular mechanisms by which spatially segregated TAM populations contribute to metastatic progression across different stages, as well as the role of the recently characterized BAMs at leptomeningeal interfaces, remain poorly understood.

*Drosophila* possess a conserved innate immune system similar to that of vertebrates, with hemocytes (plasmatocytes) representing the major cellular population of phagocytic cells, functionally analogous to mammalian macrophages (Ferrandon et al., 2007; Hoffmann, 2003). Plasmatocytes are professional phagocytes that mediate pathogen clearance through recognition and engulfment of invading microorganisms and respond to immune challenge through conserved NF-κB, MAPK and JAK-STAT signaling pathways (Charroux & Royet, 2009; Strand, 2026; Vlisidou & Wood, 2015). Like mammals, *Drosophila* plasmatocytes are also recruited to tumor sites, and are functionally analogous to mammalian TAMs (Pastor-Pareja et al., 2008). We therefore refer to *Drosophila* plasmatocytes as TAMs from here on. *Drosophila* TAMs exhibit context-dependent anti- and pro-tumorigenic functions restricting tumor growth through phagocytosis and apoptosis induction in some genetic contexts (Parisi et al., 2014; Voutyraki et al., 2023), while promoting tumor invasion and growth in other contexts (Cordero et al., 2010; Hirooka et al., 2025; Zhao et al., 2025) directly mirroring the functional duality of mammalian TAMs. Recent single-cell sequencing of *Drosophila* TAMs reveals transcriptional heterogeneity TAM populations that correspond to distinct functional states (Khalili et al., 2023; Yarikipati & Bergmann, 2026). This transcriptional continuum of multiple functional states rather than a binary classification is similar to the phenotypic plasticity of mammalian TAMs (Murray et al., 2014; Pittet et al., 2022). While several of these studies clearly establish similarities between *Drosophila* and mammalian TAMs in tumor progression at the primary site, the function of *Drosophila* TAMs at distant metastasis sites remains largely unexplored.

In this study, we utilize the adult *Drosophila* brain to investigate the cellular mechanisms by which *lgl*^-/-^ NSC-derived tumors interact with and remodel brain barrier interfaces during metastatic colonization, revealing a key role for macrophages in this process. We show that *lgl*^-/-^ tumors metastasize to the outer brain surface and progressively deform the neuronal cell cortex without infiltrating the neuropil, resembling the leptomeningeal metastasis wherein tumor cells colonize the pia matter without necessarily infiltrating the brain parenchyma (Ozair et al., 2025). This metastatic colonization occurs through a collective mode of invasion, wherein tumor cells enwrap the brain surface in a coordinated manner supported by expression of DE-cad and DN-cad and the formation of adherens junction-like structures. We further show that metastatic *lgl*^-/-^ tumors compromise both the functional and structural integrity of the BBB. Despite this, tumor cells failed to breach the SPG layer, and surprisingly, genetic ablation of the SPG alone was insufficient to promote deeper invasion, indicating the existence of additional mechanisms that restrict tumor cell entry into the neuropil.

In contrast to previously documented role of transplanted *lgl^-/-^* tumor in breaching BM by secreting MMPs in ovarian metastasis (Beaucher et al., 2007; Woodhouse et al., 1998), we found that tumor cells were unable to breach the BM, but they do result in remodeled and thickened the BM. We found that *lgl*^-/-^ tumors actively recruit TAMs to the metastatic site in a spatial pattern, and the genetic depletion of TAMs significantly reduced brain metastatic as evident by reduced tumor burden and brain deformation without affecting initial tumor cell spread or dissemination in host flies.

Overall, we put forward a novel *in vivo* model of tumor interaction with brain barrier interfaces, and establish a key role for macrophages in facilitating distant organ metastasis in *Drosophila*. Given the poorly understood mechanistic role of TAMs in human cancer metastasis, and the limited ability of mammalian models to provide cellular resolution, real-time imaging access, and fast genetic perturbation, our *Drosophila* model fills a critical gap in the field.

## RESULTS

### Tumors derived from *lgl* mutant NSCs metastasize to the adult *Drosophila* brain

Our previous work established that *lethal-giant-larvae* mutant (*lgl^-/-^*) NSC tumors transplanted into the adult fly exhibit metastatic behavior and invasive growth (Khan & Rusan, 2025). Serial transplantation of *lgl^-/-^* NSC tumors at the T3-T5 stage provides a robust metastasis model suitable for large-scale experimentation. This approach reliably results in tumor cell dissemination to the adult brain in 100% of the host flies with extensive tumor burden observed within 10-12 days post-transplantation (Figure S1A).

We first aimed to determine how metastasized *lgl^-/-^* NSC tumors (from the transplantation site) impact the brain architecture. To visualize the neuronal cell cortex, we expressed mCD8-GFP under the control of *cortex-glia-GAL4* (Kremer et al., 2017) in both controls (media injected) and *lgl^-/-^* tumor-injected flies. The neuronal cell cortex is composed of neuronal cell bodies ensheathed by cortex glia. These glial cells form a well-organized layer along the brain surface and extend inward to compartmentalize the neuronal soma (Figure 1A). Higher-magnification imaging reveals the architectural organization of this network, wherein interconnected cortex glial cells create a honeycomb-like lattice (Figure 1A1).

**Figure 1.**
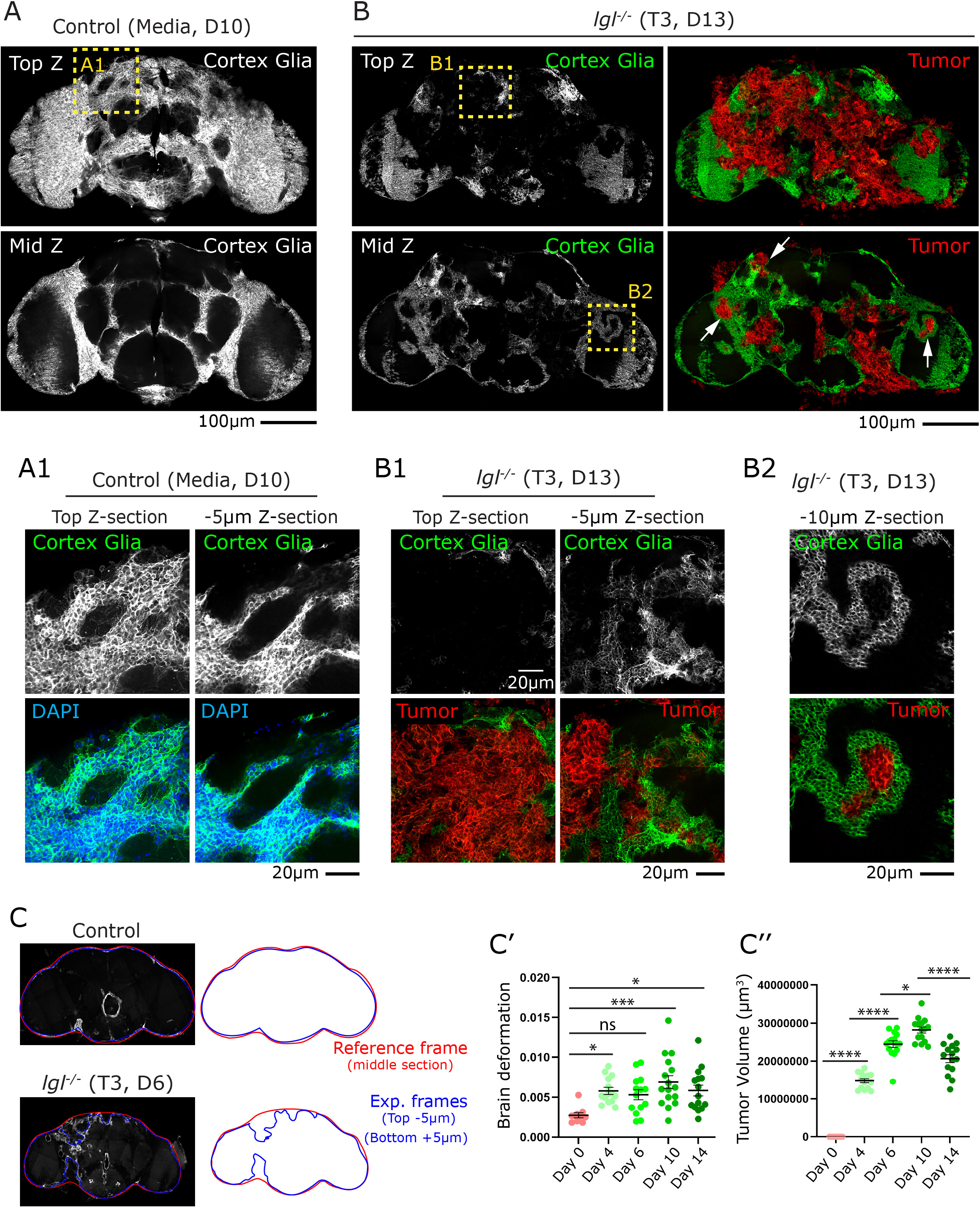
*lgl*^-/-^ tumors deform the *Drosophila* brain and disrupt neuronal cell cortex. **(A)** Representative confocal z-stack images (Top Z and Mid Z projections) of a control (Media, Day 10) *Drosophila* brain showing cortex glia (grey scale). **(A1)** High-magnification images of the boxed region in **A** showing cortex glia (green) and nuclei (DAPI, cyan) at the top and -5 µm z-sections. **(B)** Representative confocal z-stack images (Top Z and Mid Z) of an *lgl^-/-^* brain (T3, Day 13) showing cortex glia (green) and tumor cells (red). White arrows indicate sites of tumor-glia contact. **(B1)** High-magnification images of the region boxed in **B** showing cortex glia (green) and tumor (red) at the top and -5 µm z-sections. **(B2)** High-magnification image of the boxed region in **B** at the -10 µm z-section showing cortex glia (green) surrounding tumor mass (red). **(C)** Schematic illustrating the brain deformation quantification method. Brain outlines from the reference frame (middle section, red) are compared with experimental frames (top −5 µm and bottom +5 µm, blue) in a control and an *lgl^-/-^* (T3, Day 6) brain. **(C’)** Quantification of brain deformation at Day 0, 4, 6, 10, and 14 in control and *lgl^-/-^*tumor-bearing flies. **(C”)** Quantification of tumor volume (µm³) across the same time course. Data are presented as mean ± SEM, one-way ANOVA with Tukey’s multiple comparisons test was used. P < 0.05 was considered statistically significant. Significance levels are indicated as follows: ****P < 0.0001, ***P < 0.001, **P < 0.01, and *P < 0.05.

*lgl*^-/-^ tumor-injected flies typically succumb by Days 12-14 due to high tumor burden (Khan & Rusan, 2025). To examine advanced stages of brain metastasis, we dissected brains on Day 13. At this stage, the neuronal cell cortex had lost its characteristic organization, as indicated by the *apparent* absence of cortex glial cells along the brain surface (Figure 1B, Top Z). However, whole-brain imaging revealed that cortex glial cells were not eliminated; rather, the expanding tumor mass distorted the neuronal cell cortex architecture and physically displaced cortex glial cells toward the interior of the brain (Figure 1B, Mid Z; 1B1, -5 µm). In some cases, tumor cells displaced cortex glial cells into deep brain regions, giving rise to highly abnormal and complex cellular architectures (Figure 1B, Mid Z; 1B2, -10 µm). Thus, cortex glial cells remained in close contact with the tumor cells and were mechanical remodeled and displaced inward rather than lost.

To further validate these observations, we stained brains for Elav (Embryonic lethal abnormal vision) to label neuronal cell bodies. In control brains, Elav-positive neurons were neatly aligned along the brain periphery, forming a well-organized neuronal cell cortex (Figure S1B, B1; S1b, b1). In contrast, tumor-injected brains exhibited a markedly distorted neuronal cell cortex, with disrupted alignment at the brain surface (Figure S1C, C’; S1c, c1) and similar deformation extending into deeper brain regions (Figure S1C1, C1’; S1c1, c1’). Together, these findings indicate that *lgl*^-/-^ tumors successfully metastasize to the adult fly brain and induce pronounced structural deformation of the neuronal cell cortex. Consistent with our previous report (Khan & Rusan, 2025), tumor cells were not observed infiltrating the neuropil in any of the samples examined.

To quantify the extent of this deformation, we implemented a standardized brain deformation index that enables objective comparison across conditions (Figure 1C). We used the outline of the brain at two experimental positions, 5µm from the top and 5µm from the bottom (Figure 1C, Figure S2A, blue outlines), which we then compared to a reference position that outlines the middle section of the brain (Figure 1C, Figure S2A, red outline). For the tumor brains, the position of the reference outline was drawn where a normal brain would be positioned; this was done by choosing the middle section as the reference and based on our high familiarity with the highly regular wild-type brain topology (Figure 1C, S2B). All outlines were based on the position of the outermost layer of the BBB, which we labeled by expressing GFP under *perineurial-glia (PNG)-GAL4*. This PNG layer consists of a ∼1µm sheet of thin, elongated cells covering the entire brain surface (Kremer et al., 2017). Using these outlines, we determined the extent of brain deformation by calculating the mean deviation of the experimental plane from the reference plane (Figure S2C).

Analysis of brain surface deformation as a function of days post-transplantation revealed a progressive pattern of structural change over time. Metastasized tumors induced detectable deformation of the brain surface as early as Day 4 and Day 6, which increased significantly through Day 10 but declined significantly by Day 14 (Figure 1C, C’ and S3A-E & S3A’-E’). We also quantified tumor burden by measuring the overall tumor volume associated with the whole brain, our analysis revealed that tumor volume increased progressively through Day 4 to Day 10 caused by increase in tumor expansion at the brain surface as well as away from the brain surface (Figure 1C’’ and S3A’’-D’’). However, tumor volume declined significantly by Day 14 (Figure 1C’’and S3E”) similar to the decrease in brain deformation (Figure 1C’). We hypothesize that these declines at Day 14 are a result of increased lethality among flies with high tumor burden such that flies with faster tumor progression do not survive to this stage. Based on these findings, we focused all subsequent analyses on Days 10-11, a window that captured both the greatest extent of brain surface deformation and a sufficient number of surviving animals.

### *lgl^-/-^* NSC tumors metastasize to the brain show characteristics of collective invasion

Next, we aimed to determine whether the observed brain deformation results from passive adherence to the brain surface or through an active mechanism used by tumor cells. To address this, we examined the overall spatial organization and individual cell morphology of metastasized *lgl^-/-^* tumor cells at Day 10. Our analysis revealed that the tumor mass expanded along the brain surface (Figure 2A, surface view) as well as orthogonally into the brain, driving the deformation observed previously (Figure 2A, blue arrows). Three-dimensional reconstruction of confocal z-stacks (using Imaris software) revealed that tumor cells collectively enwrap the brain surface in a sheet-like manner (Video 1).

**Figure 2.**
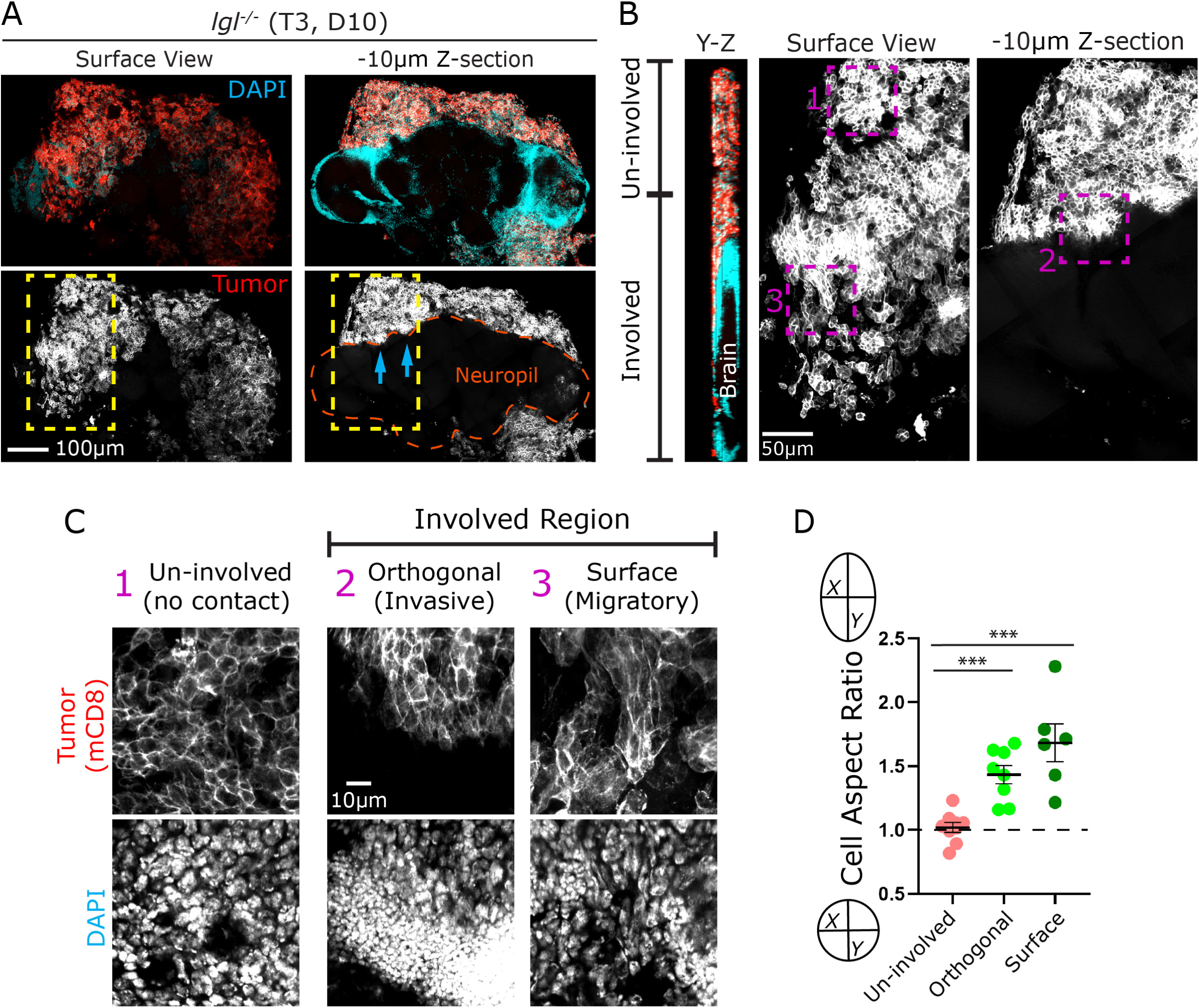
*lgl*^-/-^ deficient tumor cells adopt distinct morphologies depending on their mode of brain invasion. **(A)** Representative confocal images of an *lgl^-/-^* brain (T3, Day 10) shown at the surface view and −10 µm z-section merged channels show DAPI (cyan) and tumor cells (red). The orange dashed outline in the −10 µm z-section demarcates the neuropil region, blue arrows indicated site of brain deformation by tumor. **(B)** Orthogonal Y-Z view (left), surface view (middle), and −10 µm z-section (right) of the same brain, illustrating the distinction between un-involved (no tumor contact) and tumor-involved regions along the brain surface. **(C)** High-magnification images of the three regions indicated in **B**, showing tumor cell morphology (mCD8, grey) and nuclear morphology (DAPI, grey scale). Region **1** (un-involved, no contact) shows compact, rounded tumor cells; region **2** (orthogonal/invasive) shows tumor cells invading perpendicular to the brain surface; region **3** (surface/migratory) shows elongated tumor cells migrating along the brain surface. **(D)** Quantification of cell aspect ratio (Y/X) for tumor cells in un-involved, orthogonal (invasive), and surface (migratory) regions. Data are presented as mean ± SEM, one-way ANOVA with Dunnett’s multiple comparisons test was used. P < 0.05 was considered statistically significant. Significance levels are indicated as follows: ****P < 0.0001, ***P < 0.001, **P < 0.01, and *P < 0.05.

To understand the mode of metastatic invasion, we classified the spatial relationship between tumor cells and the brain surface into two broad categories (Figure 2B): ‘Un-involved’ regions with no direct contact with the brain surface (Box 1) and ‘Involved’ regions in direct contact with the brain (Boxes 2 and 3). Within the involved regions, we further distinguished between tumor cells oriented orthogonally to the brain surface (Box 2 at -10um) and those spreading laterally along it (Box 3 on the surface). To assess whether these spatial differences were accompanied by changes in cell morphology, we quantified the cell aspect ratio (long-axis to short-axis) across all three regions (Figure 2D). Tumor cells in un-involved regions displayed round profiles with an aspect ratio close to 1 (Figure 2C1, 2D), whereas tumor cells in both surface and orthogonal involved regions shifted toward an elongated ellipsoidal morphology, with aspect ratios approaching 1.5 (Figures 2C2-C3, 2D). This shift toward an ellipsoidal cell shape is a recognized morphological hallmark of mesenchymal motility and metastatic behavior (Friedl & Gilmour, 2009; Friedl & Wolf, 2003). The observed diversity in tumor cell morphology and spatial organization suggests a highly coordinated invasive process underlying *lgl*^-/-^ tumor metastasis to the brain. This behavior is reminiscent of collective cancer cell migration and invasion, wherein cohesive tumor clusters employ functionally specialized leader cells exhibiting mesenchymal morphology and follower cells that contribute through passive mechanical pushing to collectively navigate and invade host tissue (Friedl & Mayor, 2017; Yamamoto et al., 2023).

To test for markers of collective cancer cell migration in *lgl^-/-^* tumor, we examined the expression of E-cadherin (E-cad) and N-cadherin (N-cad), cell adhesion proteins well known for their role in mediating cell-cell contacts and promote invasive motility across several cancer types (Theveneau & Mayor, 2012; Zisis et al., 2022). In control larval brains, DE-cad localizes predominantly to the cell membrane of NSCs and their immediate progeny at sites of contact between NSCs, GMCs, and glial cells, without forming canonical adherens junctions (Figure S4A; (Almeida & Bray, 2005; Banach-Latapy et al., 2023; Doyle et al., 2017; Dumstrei et al., 2003; Fung et al., 2008). *lgl*^-/-^tumor cells that metastasized to the brain exhibited robust DE-cad expression in two distinct subcellular patterns: 1) cortical punctate labeling similar to larval NSCs (Figure 3A, blue arrow), and 2) dense plaque-like accumulations at cell-cell interfaces consistent with adherens junction-like structures (Figure 3A, magenta arrow).

**Figure 3.**
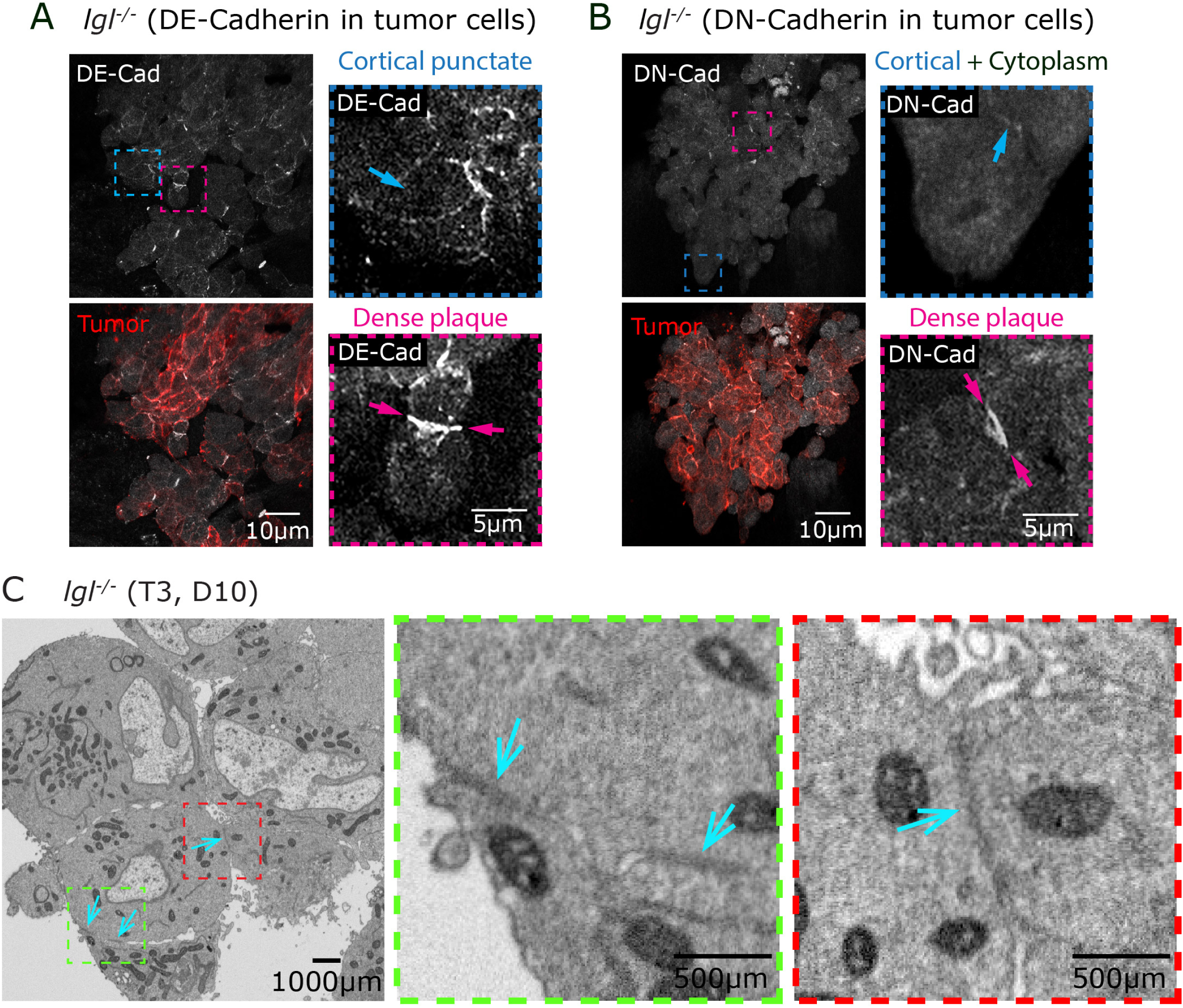
*lgl*^-/-^ deficient tumor cells display distinct cadherin localization. **(A)** Representative confocal images of *lgl^-/-^* tumor cells showing DE-Cad (grey scale) and tumor cells (red). Two magnified insets highlight distinct DE-Cad localization patterns: cortical punctate distribution (blue box, blue arrow) and dense plaque accumulation at cell-cell contacts (magenta box, magenta arrow). **(B)** Representative confocal images of *lgl^-/-^* tumor cells stained DN-Cad (grey) and tumor cells (red). Magnified insets show DN-Cad localization in both cortical and cytoplasmic compartments (blue box, blue arrow) and dense plaque formation (magenta box, magenta arrow). **(C)** Transmission electron microscopy (TEM) images of an *lgl^-/-^* brain (T3, Day 10) showing electron-dense junctions between two tumor cells cyan arrows), also magnified in green box and red box.

Unlike DE-cad, DN-cad is largely absent from NSCs and their progeny but is strongly enriched in mature neurons, where it mediates synaptic adhesion without forming canonical adherens junctions (Figure S4B; (Dumstrei et al., 2003; Fung et al., 2008; Iwai et al., 2002; Kurusu et al., 2012). Strikingly, despite their NSC origin, metastasized *lgl*^-/-^ tumor cells exhibited widespread DN-cad expression (Figure 3B). DN-cad localized in a punctate pattern at the tumor cell surface (Figure 3B, blue arrow), alongside a substantial intracellular pool, consistent with a signaling role of DN-cad in cancer cells (Mrozik et al., 2018). Furthermore, dense plaque-like DN-cad accumulations were observed at cell-cell interfaces (Figure 3B, magenta arrow), suggestive of DN-cad function in forming adherens junction-like structures. To examine the presence of cell-cell junctions more closely, we performed electron microcopy and identified distinct electron-dense junctional structures between adjacent tumor cells (Figure 3C, cyan arrows).

These results indicate that *lgl^-/-^* metastasized to the brain and form an interconnected unit that acts collectively to interact with, and deform the brain.

### Metastasized *lgl^-/-^* NCS tumor results in compromised BBB integrity and function

Despite deforming the cell cortex, *lgl^-/-^* tumor cells failed to infiltrate the neuropil, suggesting that the BBB imposes a physical or biochemical constraint. To investigate this, we sought to identify any functional or structural defects in the BBB of brains with metastatic tumors. The *Drosophila* BBB consists of a basement membrane (BM) followed by perineurial glia (PNG) and sub-perineurial glia (SPG) cell layers. Like the mammalian BBB, it is characterized by septate junctions that prevent paracellular transport of solutes and cells (Contreras & Klambt, 2023; Limmer et al., 2014).

To evaluate the functional integrity of the BBB, we utilized a 10 KDa Texas Red-conjugated Dextran exclusion assay. Control brains showed no evidence of BBB leakage, as indicated by the absence of dextran signal (Figures S5A and D). Conversely, brains in which the BBB had been genetically ablated exhibited significant dextran uptake, serving as a positive control (Figures S5B and D). Notably, brains harboring *lgl^-/-^* tumors also displayed significant dextran uptake as early as Day 6 (Figures S5C and D). This suggests that the metastasized tumor compromised BBB integrity, potentially through direct mechanical disruption of glial layers or via indirect inflammatory signaling, as previously documented in other *Drosophila* tumor models (Kim et al., 2021).

To analyze the integrity of BBB cell layers we examined PNGs at the tumor-brain interface by expressing mCD8-GFP under a PNG-specific Gal4 driver. As previously established, control PNG cells exhibit a characteristic morphology of thin, elongated, and interdigitated cells with overlapping membrane ledges (Figure 4A, A1, A1’; red arrow) (Kremer et al., 2017). The control brains exhibited normal tiling of PNG (Figure 4C, control brain). We then examined PNG cells in two regions of brains with metastasized *lgl^-/-^* tumors: ‘tumor involved’ regions and ‘no tumor’ regions (an internal control). Our analysis revealed that the PNG layer in involved regions incurred two significant defects: 1) Physical disruption of the PNG, either loss of cellular overlap (Figure 4B, B1, B1’, red arrow), or complete loss of PNG cells (Figure 4B, B2, red arrow). 2) Intracellular granule accumulation inside the PNG cells (Figure 4B3). Quantification across multiple regions of interest (ROIs) demonstrated that the loss of PNG-PNG contact was predominantly associated with the involved region, and was significantly low in both “no tumor” region and control brains (Figure 4C). However, the granular phenotype was observed in PNG cells throughout the brain with metastasized tumors, irrespective of direct contact with the tumor; in contrast, granules were completely absent in control brains (Figure 4D).

**Figure 4.**
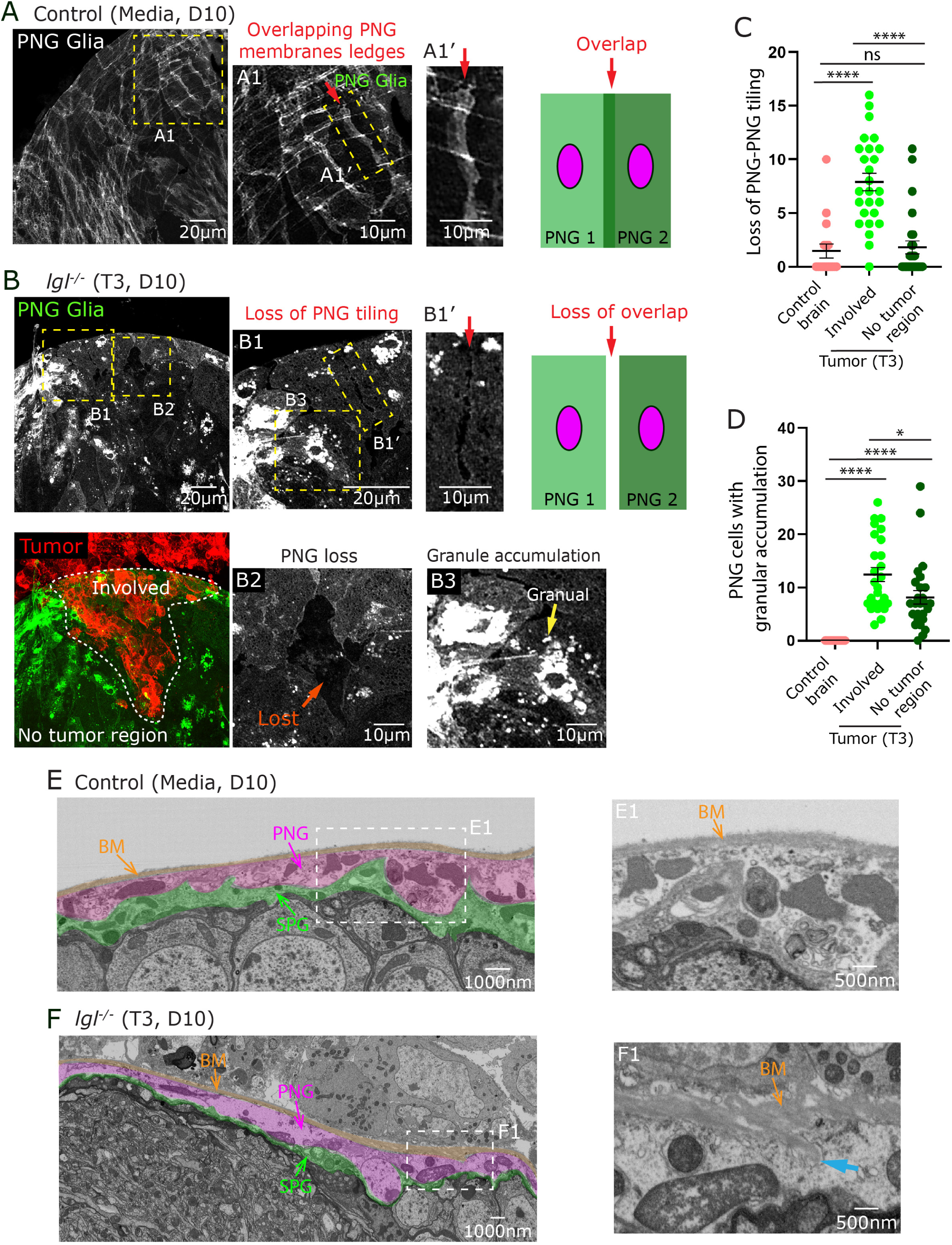
*lgl^-/-^* tumors disrupt ultrastructural organization of the BBB. **(A)** Representative confocal image of a control (Media, Day 10) brain showing PNG Glia (grey scale). **(A1)** High-magnification image showing overlapping PNG membrane ledges (red arrow). **(A1′)** High-magnification image of the PNG-PNG membrane overlap (red arrow). Schematic illustrates the normal tiling arrangement in which adjacent PNG cells (PNG1 and PNG2) (membrane overlap, red arrow). **(B)** Representative confocal images of an *lgl^-/-^* brain (T3, Day 10). **(B1)** High-magnification image showing loss of PNG-PNG tiling in the tumor-involved region. **(B1′)** High-magnification image showing loss of membrane overlap between adjacent PNG cells (red arrow). Schematic illustrates the disrupted tiling between two adjacent PNG1 and PNG2 (loss of overlap, red arrow). **(B2)** High-magnification image showing complete absence of PNG cells in the tumor-involved region (orange arrow). **(B3)** High-magnification image showing intracellular granule accumulation within PNG cells in the tumor-proximal region (yellow arrow). **(C)** Quantification of PNG-PNG tiling loss events per brain in control brains, tumor-involved regions, and no-tumor regions. **(D)** Quantification of the number of PNG cells displaying granular accumulation per brain across the same three conditions. **(E)** FIB-SEM overview image of a control (Media, Day 10) brain showing the normal ultrastructural organization of the glial BBB. The basement membrane (BM, orange overlay), perineurial glia (PNG, magenta overlay), and subperineurial glia (SPG, green overlay) are indicated. **(E1)** High-magnification image showing the intact BM (orange arrow) in the control condition. **(F)** FIB-SEM overview image of an *lgl^-/-^* brain (T3, Day 10) showing tumor cells pushing the brain surface. **(F1)** High-magnification image showing BM thickening (orange arrow), BM pinching or accumulation in PNG (blue arrow). Data are presented as mean ± SEM, one-way ANOVA with Tukey’s multiple comparisons test was used. P < 0.05 was considered statistically significant. Significance levels are indicated as follows: ****P < 0.0001, ***P < 0.001, **P < 0.01, and *P < 0.05.

The observation that tumor cells physically disrupted the structural integrity of PNG cell layer where tumor cell made direct contact with the brain suggested that a specific interaction between tumor cells and BBB at the metastatic sites. To further validate this, we performed Focused Ion Beam-Scanning Electron Microscopy (FIB-SEM) to visualize the ultrastructural relationship between tumor cells and the BBB layers at nanometer resolution. FIB-SEM imaging of control brains revealed the characteristic tri-layered BBB architecture comprising a thin, uniform basement membrane (BM, orange overlay) followed by the PNG (magenta overlay) and SPG (green overlay) cell layers (Figure 4E and S6A). In tumor-injected flies, FIB-SEM confirmed that tumor cells exert direct mechanical pressure on the brain surface at sites of contact, producing pronounced deformation of the underlying PNG cells and flattening of the SPG layer (Figure 4F and S6B). Consistent with our confocal data, loss of PNG-PNG contact was evident at these sites (Figure S6B-B1, magenta arrow) and not in controls (Figure S6A-A1, magenta arrows). Despite the structural deformation of the PNG cells, the BM remained intact at all tumor contact sites examined, with no evidence of BM breach (Figure 4F). Strikingly, however, the BM at some sites of direct tumor contact displayed pronounced ruffling and was markedly thickened, measuring approximately 500-600 nm compared to 200-300 nm in control brains (Figure 4E1, F1, orange arrows). This suggests active remodeling of the BM in response to mechanical pressure from the tumor cells. Intriguingly, we also observed ECM-filled vesicles within PNG cells in continuity with BM, indicating a PNG role in BM remodeling (Figure 4F1 and S6 B2 and B3, blue arrows).

FIB-SEM data revealed pronounced flattening of the SPG layer at sites of direct tumor contact (Figure 4F, green overlay), prompting us to assess whether this mechanical deformation compromised SPG barrier integrity. To test this, we expressed mCD8-GFP under an SPG-specific Gal4 driver to visualize SPG cell morphology and cell-cell contacts (Kremer et al., 2017). In control brains, SPG cells form large, thin, flattened sheets tightly sealed by septate junctions between neighboring cells (Figure 5A). In tumor-bearing brains, SPG cells at involved regions were displaced inward by the advancing tumor mass; however, SPG-SPG contacts remained largely intact in the vast majority of cases (>95%), as visualized by continuous membrane overlap (Figure 5B and B1). Only rarely (<5%) did we observe a detectable loss of SPG-SPG contact (Figure 5B and B2). These findings demonstrate that despite pronounced deformation of the brain surface and structural disruption of the PNG cell layer, *lgl^-/-^* tumor cells are largely unable to breach the SPG barrier.

**Figure 5.**
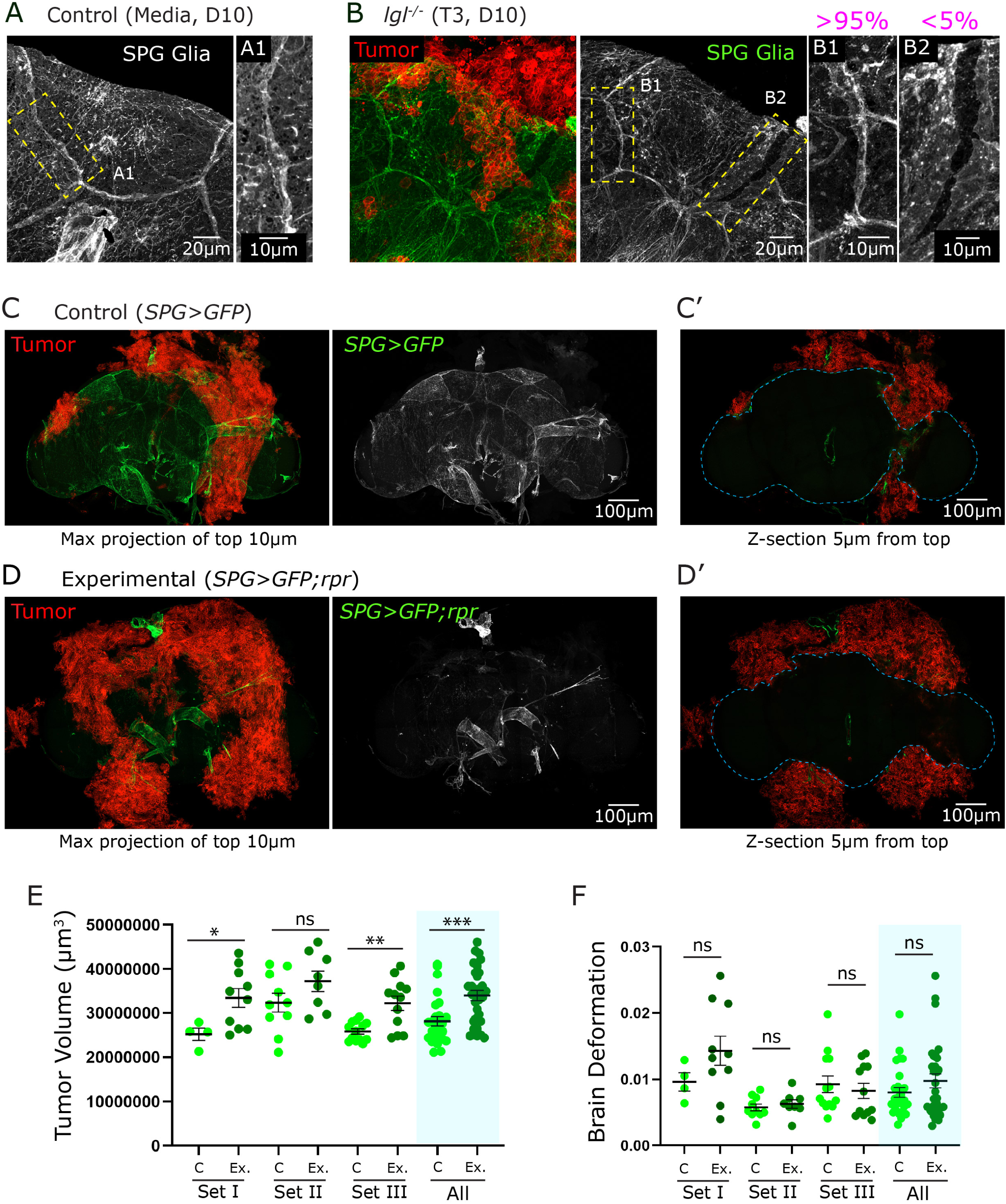
SPG ablation does not significantly alter brain deformation but modestly affects tumor volume. **(A)** Representative image of a control (Media, Day 10) brain showing SPG Glia (grey scale). **(A1)** High-magnification image of the boxed region showing the intact SPG morphology. **(B)** Representative images of an *lgl^-/-^* brain (T3, Day 10) showing tumor cells (red) and SPG Glia (grey scale). **(B1)** High-magnification image of the SPG region with greater than 95% SPG coverage retained in the tumor-involved area (>95%). **(B2)** High-magnification image showing loss of SPG-SPG contact at the tumor-brain interface (<5%). **(C)** Representative images of a control (*SPG>GFP*) brain with transplanted tumor cells, merged maximum projection of the top 10 µm showing tumor (red) and SPG>GFP signal (green). **(C’)** Single z-section 5 µm from the top showing tumor (red) and the brain boundary (cyan dashed line). **(D)** Representative images of an experimental (*SPG>GFP;rpr*) brain in which SPG cells are genetically ablated via co-expression of the pro-apoptotic gene *rpr*, merged maximum projection of the top 10 µm showing tumor (red) and residual SPG>GFP signal (green) following ablation. **(D’)** Single z-section 5 µm from the top showing tumor (red) and brain boundary (cyan dashed line) in the SPG-ablated condition. **(E)** Quantification of tumor volume (µm³) in control (C) and experimental (Ex.) animals across Sets I, II, and III and pooled across all sets (blue shaded region, All). **(F)** Quantification of brain deformation in control (*SPG>GFP*, C) and experimental (*SPG>GFP;rpr*, Ex.) animals across three independent experimental sets (Set I, Set II, Set III) using -5 µm top and reference frame and pooled across all sets (blue shaded region, All). Data in **E** and **F** are presented as mean ± SEM with individual data points shown. Two-tailed Welch’s t-test was used, \**P* < 0.05 is considered significant. Significance levels are indicated as follows: ****P < 0.0001, ***P < 0.001, **P < 0.01, and *P < 0.05.

Together with the intact BM observed by FIB-SEM, these data indicate that the SPG layer constitutes a second line of defense that, in concert with the BM, restricts tumor cell access to deeper brain regions, and prevents parenchymal infiltration.

### The SPG layer does not serve as the primary physical barrier to *lgl^-/-^* NSC tumor invasion

To directly test if the SPG layer functions as the principal barrier preventing tumor cells from infiltrating into deeper brain regions, we genetically ablated SPG cells by expressing the pro-apoptotic gene reaper (*rpr*) in adult flies using an SPG-specific Gal4 driver combined with a temperature-sensitive Gal80^ts^ system (methods). This manipulation led to robust loss of SPG cells in the adult fly brain. In contrast, control flies lacking *rpr* expression retained an SPG cell layer with intact SPG-SPG connection (Figure 5C-D).

Interestingly, SPG ablation led to a visible increase in overall tumor burden across all three independent paired datasets (Figure 5C, D). We quantified this by measuring tumor volume across all sets, which consistently showed that loss of SPG enhanced the growth of metastatic *lgl^-/-^* tumors (Figure 5C, D, E). Surprisingly despite an increase in tumor burden, SPG loss did not enhance brain deformation, nor did we detect evidence of *lgl^-/-^* tumor cells in the neuropil region of SPG-deficient brains (Figure 5C’, D’; 5F). Together, these results indicate that the SPG layer does not act as the sole barrier to infiltration of *lgl^-/-^* tumors in to the brain parenchyma. The persistence of a BM, although remodeled, at tumor contact sites may account for the lack of change in brain deformation and lack of tumor cell infiltration following SPG removal, suggesting that the BM rather than the SPG constitutes the primary physical barrier to parenchymal infiltration. The increase in tumor volume observed in SPG-ablated brains might reflects heightened exposure to brain-derived growth factors or elevated stress-related signaling, creating a locally permissive microenvironment for peripheral tumor growth and expansion without any significant effect on brain deformation.

### Metastasized *lgl^-/-^* NCS tumor recruits tumor-associated-macrophages (TAMs)

The structural disruption of the PNG and active remodeling of the BM at tumor contact sites raised the question of whether the innate immune system responds to, and participates in, this process. Given that *Drosophila* macrophages are known to respond to tissue damage and BM disruption (Pastor-Pareja et al., 2008), and that our data revealed remodeling of BM at tumor contact sites, we hypothesized that macrophages could actively be recruited and contribute to tumor-brain interactions.

Tissue-resident macrophages in adult flies exist as sedentary immune cells beneath the cuticle wall and within the head capsule (Bosch et al., 2019). In control brains injected with media, macrophages were primarily localized near the brain surface and respiratory epithelium (trachea), as evident by anti-Nimrod C1/2 (NimC) staining (Figure 6A). The majority of macrophages were organized in discrete clusters and exhibited a rounded morphology of approximately 10-15 µm in diameter, consistent with a quiescent, non-activated state (Figure 6A1, A2). These observations were further confirmed using the *srpHemo-3xmCherry* reporter line (Gyoergy et al., 2018), which similarly revealed an absence of activated or phagocytic macrophages under control conditions (Figure S7A; S7A1, A2). In contrast, brains bearing metastatic *lgl*^-/-^ tumors exhibited a marked reorganization of the macrophage population, characterized by the loss of discrete clusters typical of quiescent cells. Instead, the majority of macrophages appeared mobilized and recruited to the metastatic tumor mass as tumor-associated macrophages (TAMs) (Figure 6B). Higher-magnification analysis revealed pronounced morphological changes in TAMs, including an approximately 5-6-fold increase in cell size and an elongated morphology consistent with a activated phenotype (Figure 6B1 and 6B2). Consistent with these findings, the *srpHemo-3xmCherry* reporter line revealed similar morphological changes in TAMs, including characteristics typical of activated macrophages exhibiting elongated cell morphology (Figure S7B).

**Figure 6.**
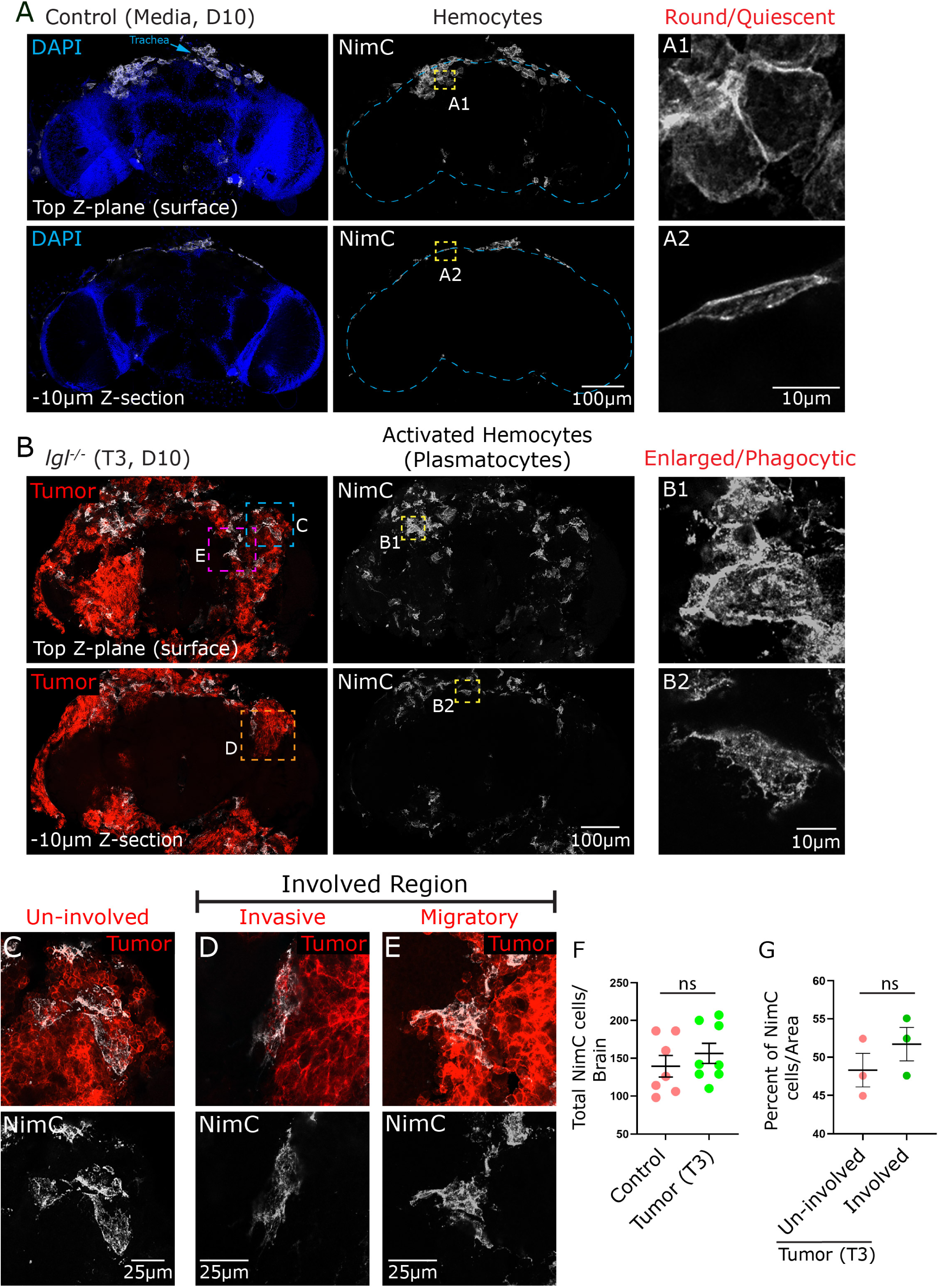
*lgl^-/-^* tumors recruit activated macrophage in a distinct spatial pattern. **(A)** Representative images of a control (Media, Day 10) brain at the top z-plane (surface) and −10 µm z-section. Left column shows DAPI (blue) with trachea indicated (cyan arrow). Middle column shows NimC-positive hemocytes (grey scale) with brain boundary (cyan dashed line). **(A1-A2)** High-magnification images of showing round/quiescent hemocyte morphology in control brains. **(B)** Representative images of an *lgl^-/-^* brain (T3, Day 10) at the top z-plane (surface) and −10 µm z-section. Middle column shows NimC-positive hemocytes (grey scale). **(B1-B2)** High-magnification images showing enlarged/phagocytic hemocyte (activated plasmatocytes) in tumor-bearing brains. **(C)** High-magnification images of the un-involved (no tumor contact) region indicated in **B** (blue box). **(D)** High-magnification images of the invasive region of the involved area indicated in **B** (orange box). **(E)** High-magnification images of the migratory region of the involved area indicated in **B** (magenta box). **(F)** Quantification of total NimC-positive hemocyte number per brain in control and *lgl^-/-^* tumor. **(G)** Quantification of the percentage of NimC-positive cells per unit area in un-involved versus tumor-involved regions across whole brain, n=3. Data in **F** and **G** are presented as mean ± SEM with individual data points shown. Two-tailed Welch’s t-test was used, \**P* < 0.05 is considered significant. Significance levels are indicated as follows: ****P < 0.0001, ***P < 0.001, **P < 0.01, and *P < 0.05.

Next, we investigated the spatial distribution of TAMs across the metastatic tumor mass. Analysis of multiple regions of interest (ROIs) revealed that TAMs were not exclusively confined to tumor cells directly engaging the brain surface; they were also recruited to tumor cells in un-involved regions (Figure 6C). Within involved regions, TAMs were frequently observed at the leading edges of tumor cells oriented orthogonally to the brain surface (Figure 6D), as well as at the leading edges of tumor cells spreading laterally along the brain surface (Figure 6E). Quantification of NimC-positive cells revealed no significant difference in total TAM numbers between control and tumor-injected brains (Figure 6F), suggesting a dynamic spatial redistribution rather than proliferative expansion of the macrophage population. Additionally, quantification of TAMs in un-involved versus involved regions indicated a slight but non-significant enrichment of TAMs in involved regions (Figure 6G). This distribution is consistent with observations in mammalian metastatic niches, where TAMs exhibit accumulating at both the invasive tumor edges and within the tumor core, a so called compartment-dependent distribution (Friebel et al., 2020; Gan et al., 2024; Karimi et al., 2023).

### TAMs facilitate metastasis of *lgl^-/-^* NCS tumor to the brain

To directly assess TAMs functions in limiting or promoting metastasis of *lgl^-/-^*tumors to the brain, we genetically ablated macrophages in adult flies using a Gal80^ts^ system combined with the expression of the pro-apoptotic genes *rpr* and *hid*. Our experimental approach significantly reduced macrophage numbers as confirmed by GFP signal and NimC staining compared to the controls (Figure 7A,B). Strikingly, macrophage ablation led to a significant reduction in tumor burden in the brain as measured by tumor volume across two independent experiments (Figure 7A-C). This reduction in tumor burden was also accompanied by a corresponding decrease in brain deformation, further indicating diminished metastatic engagement of tumor cells with the brain surface (Figure 7A’-B’ and D).

**Figure 7.**
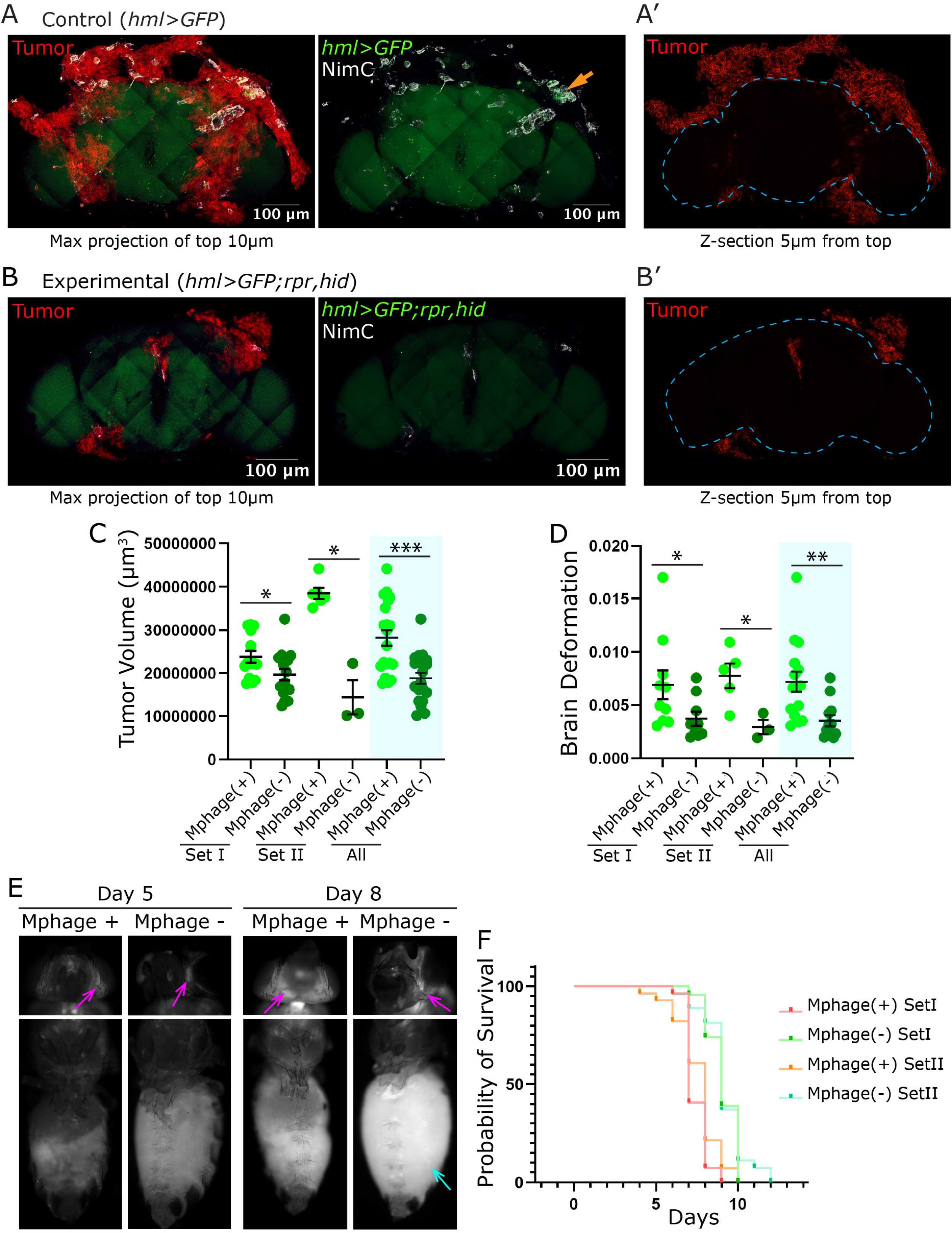
TAMs ablation reduces metastasis to brain. **(A-A’)** Representative confocal images of a control (*hml>GFP*) brain with transplanted tumor cells. **(A)** Maximum projection of the top 10 µm showing tumor cells (red) and merged *hml>GFP* hemocytes (green) with NimC staining (grey scale), orange arrow indicates a NimC-positive macrophages. (**A’)** Single z-section 5 µm from the top showing tumor cells (red) and brain boundary (cyan dashed line). **(B-B’)** Representative confocal images of an experimental (*hml>GFP;rpr,hid*) brain in which hemocytes are genetically ablated via co-expression of the pro-apoptotic genes *rpr* and *hid* under the *hml*-Gal4 driver. **(B)** Maximum projection of the top 10 µm showing tumor cells (red) and merged *hml>GFP;rpr,hid* signal (green) with NimC staining (grey scale). **(B’)** Single z-section 5 µm from the top showing tumor cells (red) and brain boundary (cyan dashed line) in the hemocyte-ablated condition. **(C)** Quantification of tumor volume (µm³) in hemocyte-present [Mphage(+)] and hemocyte-ablated [Mphage(−)] animals across two independent experimental sets (Set I, Set II) and pooled across both sets (blue shaded region, All). **(D)** Quantification of brain deformation in Mphage(+) and Mphage(−) animals across Set I, Set II, using -5 µm top and reference frame and pooled (blue shaded region, All). **(E)** Representative brightfield images of whole flies at 2X magnification from Mphage(+) and Mphage(−) conditions at Day 5 (left) and Day 8 (right) post-transplantation. Heads from the same host imaged at 5X magnification, magenta arrows indicate tumor mass visible through the cuticle. Cyan arrow indicates tumor growth in host abdomen at Day 8 in the Mphage(−) condition. **(F)** Kaplan-Meier survival curves for Mphage(+) and Mphage(−) animals across Set I (pink/green) and Set II (orange/cyan) over the course of 15 days post-transplantation. Data in **C** and **D** are presented as mean ± SEM with individual data points shown. Two-tailed Welch’s t-test was used, \**P* < 0.05 is considered significant. Significance levels are indicated as follows: ****P < 0.0001, ***P < 0.001, **P < 0.01, and *P < 0.05.

To determine whether reduced brain metastasis reflected a general impairment in tumor dissemination or a specific defect in brain colonization, we tracked tumor cells in host flies at Days 5 and 8 post-transplantation. In both control and macrophage-ablated flies, tumor cells disseminated to the head capsule by Day 5 (Figure 7E, magenta arrow), indicating that macrophage ablation does not impair the initial ability of tumor cells to spread from the abdominal injection site. By Day 8, however, both control and macrophage ablated flies showed comparable tumor signal in the head capsule despite extensive tumor growth in the abdomen of flies lacking macrophages (Figure 7E, cyan arrow). These results revealed that macrophages are dispensable for the initial spread of tumor cells to the brain and tumor growth in the head capsule. Limited brain metastatic colonization (tumor volume + brain deformation) (Figure 7A-D), suggest a role of macrophages in efficient engagement of tumor cells with the brain surface. The increased abdominal tumor growth in macrophage-ablated flies likely reflects suppression of anti-tumor immune responses (Parisi et al., 2014; Voutyraki et al., 2023). Notably, despite this extensive tumor growth, macrophage-ablated flies survived 10-20% longer than controls (Figure 7F), a delay we attribute to reduced brain invasion and diminished inflammatory signaling. Overall, our results support the idea that macrophages plays a critical function in metastatic colonization, potentially by supporting efficient expansion and engagement of *lgl^-/-^* tumors to the brain surface. The coincidence of TAMs at the leading edges of the tumor-brain interface (Figure 6B, D and E) suggests that a specialized subset of TAMs could be involved in facilitating directed migration of tumor cells along and into the brain surface. Additionally, TAMs association within the tumor core may provide pro-proliferative or pro-survival cues to support the growth of tumor cells that finally result in successful expansion and colonization of the brain.

### Potential role of TAMs in *lgl^-/-^* tumor metastasis through BM remodeling at the brain surface

To gain deeper insight into the cellular interaction between the tumor, TAMs and the brain surface, we performed FIB-SEM imaging of tumor-brain interface. FIB-SEM revealed that TAMs not only associated closely with tumor cells but also formed direct physical contacts between tumor cells and the brain surface (Figure 8A, green overlay), positioning themselves at the interface between the tumor and the underlying BM (Figure 8A1, green arrow).

**Figure 8.**
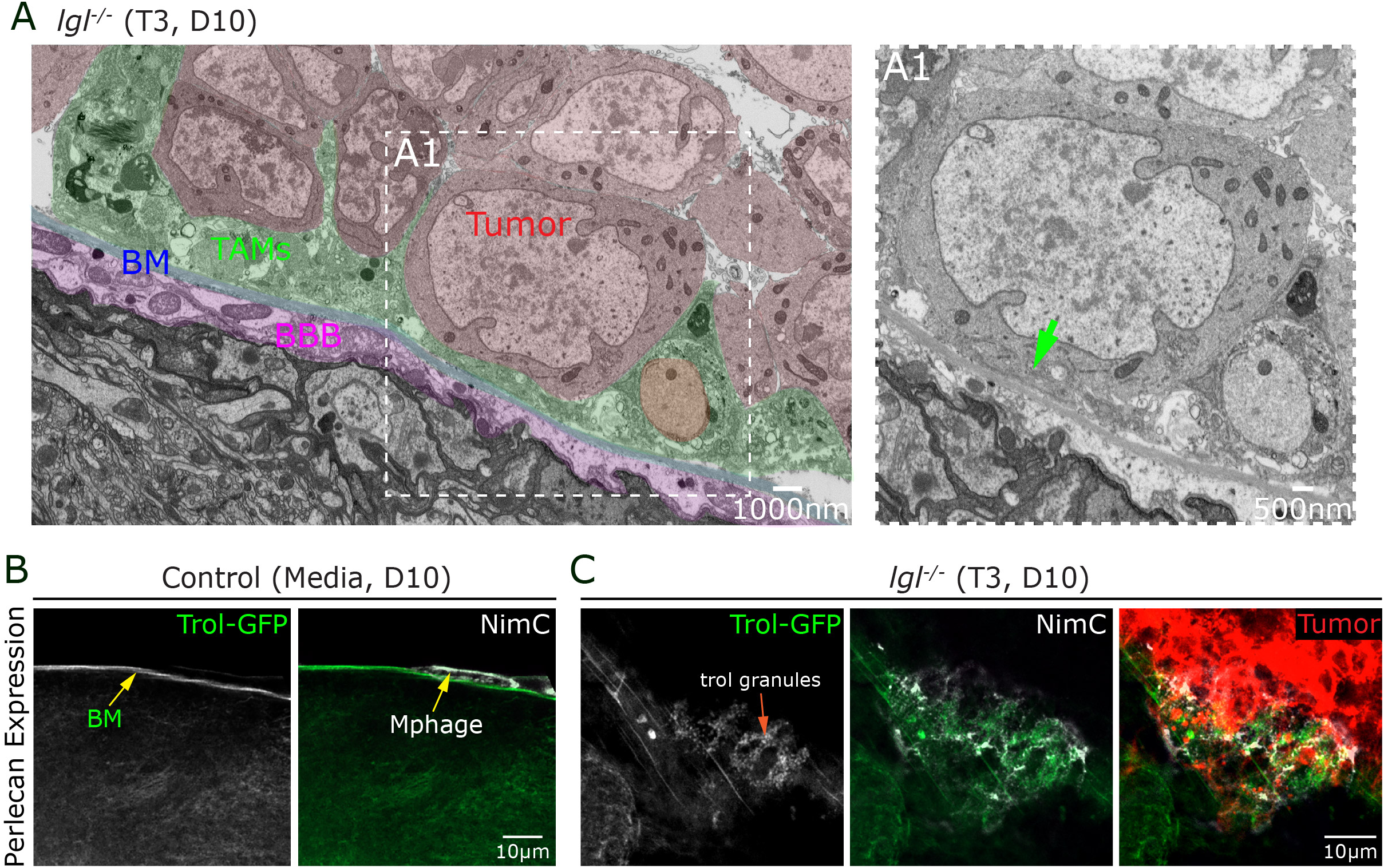
Potential role of TAMs in basement membrane remodeling at the tumor-brain interface. **(A)** False-colored FIB-SEM overview image of an *lgl^-/-^* brain (Transplant 3, Day 10) showing the spatial relationship between key cellular and extracellular components at the tumor-brain interface. The basement membrane (BM, blue overlay), Blood-brain barrier (BBB, magenta overlay), Tumor-associated macrophages (TAMs, green overlay) and tumor cells (tumor, red overlay). **(A1)** High-magnification FIB-SEM image of the boxed region in A showing ultrastructural details at the TAMs-BM-tumor interface. The green arrow indicates TAMs interfacing BM and tumor cell. **(B)** Representative confocal images of a control (Media, Day 10) brain showing Perlecan expression via the endogenous *trol-GFP*. *Trol-GFP* showing Perlecan localization along the BM (green, yellow arrow) and NimC staining showing hemocytes (Mphage, yellow arrow) distribution along the BM in control brains. **(C)** Representative confocal images of an *lgl^-/-^* brain (Transplant 3, Day 10) showing Perlecan/Trol expression in the presence of tumor. *Trol-GFP* signal (grey scale) showing ectopic Perlecan accumulation as discrete intracellular granules (trol granules, orange arrow) in the NimC positive TAMs.

TAMs are known to actively remodel the BM through enzymatic modification of ECM and by secretion of matrix proteins laying a permissive track along for tumor cells directed migration (Afik et al., 2016; Bahr et al., 2022; Sharma et al., 2012). Spatial configuration seen in our model led us to hypothesize that TAMs act as cellular intermediaries that physically bridge advancing tumor cells to the BM at the brain surface, enabling guided migration and invasion. To test the role of TAMs in BM remodeling, we examined whether TAMs express ECM components using a *Trol-GFP* to track the heparan sulfate proteoglycan Perlecan. We found that in control flies *Trol-GFP* localized exclusively to the BM with no detectable signal in hemocytes (Figure 8B and Figure S8A). Strikingly, in tumor-bearing brains, approximately 40% of TAMs contained *Trol-GFP*-positive granules, indicating active uptake or secretion of Perlecan by TAMs (Figure 8B and S8B-C). However, TAMs positive for *Trol-GFP* were found both at invasion sites coinciding with regions of BM damage and associated with tumor cells in un-involved regions away from the brain surface, with no significant spatial preference at either the invasive front or the tumor core (Figure S8B-D). Suggesting, a dual role of TAMs at tumor-BM interface and within the tumor core via BM remodeling and matrix proteins modification.

## DISCUSSION

In this study, we utilized the adult *Drosophila* brain as a genetically tractable *in vivo* platform to investigate the cellular mechanisms underlying brain metastatic colonization. Using serial transplantation of *lgl*^-/-^ NSC-derived tumors, we demonstrate that tumor cells colonize the brain surface and deform neuronal cell cortex. We show that *lgl*^-/-^tumor cells utilize collective mode of invasion to spread in a sheet-like manner and colonize the brain. Despite the structural compromise of the BBB, including disruption of the PNG cell layer at sites of direct tumor contact, tumor cells consistently failed to breach the BM and SPG cells layer, and fail to infiltrate the neuropil. Surprisingly, genetic ablation of the SPG cell layer slightly increased tumor burden without promoting parenchymal infiltration, revealing that BM potentially act a critical barrier in restricting tumor cell entry into the brain parenchyma. We further show that *lgl*^-/-^ tumors recruit macrophages (TAMs) at the tumor leading edges in contact with the brain as well as within the tumor core. Importantly, genetic depletion of TAMs significantly reduced the metastasis, as evidenced by less tumor burden and brain deformation. Interestingly, our data support TAMs function in tumor cell engagement via BM remodeling. Altogether, based on these findings we propose a model wherein TAMs facilitate metastatic colonization of the brain through two complementary functions: actively remodeling the BM through matrix protein secretion to enable guided tumor cell engagement with the brain surface, and by supporting the expansion of successfully colonized tumor cells.

### Adult *Drosophila* brain model for BBB-tumor interaction

A central observation in this study is that metastatic *lgl*^-/-^ tumors colonize the outer surface of the adult *Drosophila* brain and deform the neuronal cell cortex. This results in compromising the PNG cell layer without breaching the BM and SPG cell layer, or infiltrating the neuropil. This pattern of surface-restricted colonization bears a striking anatomical resemblance to leptomeningeal disease (LMD) in humans, where tumor cells spread through the CSF compartments and pia matter without necessarily infiltrating the brain parenchyma (Freret & Boire, 2024; Ozair et al., 2025). LMD remains one of the most poorly understood and clinically devastating forms of brain metastasis, with limited tractable *in vivo* models for mechanistic dissection (Ozair et al., 2025). The fact that *lgl*^-/-^ tumors colonize the space between the hemolymph and BM, deform the brain surface, and compromise barrier integrity without crossing the BBB. This suggest that *Drosophila* can be an important model of leptomeningeal colonization, offering a analogous system for studying tumor cells interaction with pia matter and underlying BM of the glia limitans layer (Freret & Boire, 2024; Ozair et al., 2025).

One unexpected finding was that genetic ablation of the SPG layer, the primary paracellular barrier of the *Drosophila* BBB, increased tumor burden without an evident increase in brain deformation and parenchymal infiltration. This result demonstrates that the SPG is not the principal physical barrier preventing tumor entry into the brain parenchyma, and instead implicates additional, organ-intrinsic or organ-responsive mechanisms that independently restrict tumor colonization. One candidate is the BM itself. Our FIB-SEM data revealed pronounced thickening and ruffling of the BM at tumor contact sites increasing from approximately 200-300 nm in controls to 500-600 nm in tumor-bearing brains suggestive of active ECM remodeling at the invasion interface. This remodeling likely involves contributions from both macrophages, which are enriched at the invasive front and are well-established sources of ECM-modifying enzymes, and the underlying PNG cells, which we observed to contain ECM-filled vesicles at tumor contact sites (Figure 4). Importantly, a thickened and remodeled BM is not merely a barrier in mammalian tumors, increased ECM stiffness and density are well documented to promote tumor cell proliferation, survival, and migration by activating integrin-mediated mechanosignaling and providing an enriched substrate for cell adhesion and motility (Levental et al., 2009; Paszek et al., 2005; Pickup et al., 2014). The BM thickening observed at invasion sites may therefore create a permissive microenvironment that supports tumor expansion along the brain surface. This can also help explain why SPG loss, which further exposes the tumor to brain-derived signals, increases tumor burden without enabling parenchymal infiltration. This model can be directly tested in *Drosophila* by selectively depleting specific ECM components such as Collagen IV or Laminin from macrophages or PNG cells, or by targeting enzymes that regulate BM assembly and crosslinking, such as peroxidasin or matrix metalloproteinases.

A further observation with important mechanistic implications is the granule accumulation phenotype in PNG cells throughout tumor-bearing brains, including in cells not in direct contact with the tumor. This non-cell-autonomous changes within PNG cells suggests that metastatic *lgl*^-/-^ tumors influence the BBB not solely through direct mechanical contact but also through humoral or paracrine signals that propagate across the barrier (Figure 4). This is reminiscent of how systemic cytokine signaling including interleukin and TNF-family ligands compromises BBB integrity in mammalian brain metastasis, often preceding and facilitating tumor extravasation (Arvanitis et al., 2020; Gril et al., 2018; Soto et al., 2014). In *Drosophila*, inflammatory signaling through JAK-STAT and JNK pathways is well characterized downstream of tumor-derived signals (Bilder et al., 2021; Kim et al., 2021), and the granule phenotype may reflect activation of such pathways in PNG cells responding to tumor-secreted factors. The nature of these granules and the identity of the upstream tumor-derived signals would important in uncovering BBB response to tumor metastasis.

### *Drosophila* model for investigating macrophage function in tumor metastasis

One of the key findings of this study is that genetic depletion of TAMs significantly reduces metastasis to the adult brain, demonstrating the role for innate immune cells at a distant metastatic site in *Drosophila* (Figure 7). This finding extends the functions of *Drosophila* macrophages previously documented at primary tumor sites (Cordero et al., 2010; Hirooka et al., 2025; Pastor-Pareja et al., 2008; Zhao et al., 2025) to the context of distant organ metastasis. Importantly, because depletion of TAMs reduced tumor burden and overall brain deformation without affecting tumor cell spread to the head capsule. The apparent decrease in metastasis could be explained by decrease in tumor cell engagement (physical attachment) with the brain and reduced expansion of tumor cells and eventual colonization. In support of this model we find that TAMs localize at the interface between tumor cells and outermost BM layer of the brain (Figure 8). We surmise that TAMs could play a key role in BM remodeling via matrix protein modification and secretion that would in turn be required for directed migration of tumor cells, a well-known function of TAMs in mammalian models (Afik et al., 2016; Bahr et al., 2022; Sharma et al., 2012). Directly testing this by selective depletion of matrix protein or matrix remodeling enzymes in TAMs will shed light on potential mechanism utilized by TAMs in organ metastasis.

How TAMs mechanistically support *lgl^-/-^* tumor metastasis at the brain surface remains an open question. Several functional models are plausible based on what is known from mammalian systems and *Drosophila* primary tumor biology. We show that in addition to the tumor core, TAMs are also prominently recruited at the leading edges of tumor cells engaging with the brain surface orthogonally and laterally (Figure 6). At these sites TAMs may facilitate tumor cell protrusion and motility through paracrine cytokine loops analogous to the CSF-1/EGF signaling axis described in mammary tumor invasion (Goswami et al., 2005; Wyckoff et al., 2004). Beyond the leading edges, TAMs associated with tumor cells in un-involved regions may support early colonization by suppressing anti-tumor immunity or providing trophic support, mirroring the role of macrophages in the tumor core during mammalian brain metastasis (Friebel et al., 2020; Klemm et al., 2020).

A complementary possibility is that TAMs facilitate tumor invasion by promoting an epithelial-to-mesenchymal transition (EMT)-like program in tumor cells at the leading edge. TAM-driven EMT has been documented in multiple mammalian cancer types, mediated by TGF-β, TNF, and EGF signaling that collectively downregulate epithelial adhesion molecules and upregulate mesenchymal motility programs (Bonde et al., 2012; Li et al., 2022; Su et al., 2014). The observation that *lgl^-/-^* tumor cells at the leading edges adopt an elongated, ellipsoidal morphology with aspect ratios approaching 1.5 a recognized hallmark of mesenchymal motility raises the possibility that contact with or signaling from TAMs at the tumor-brain interface induces or stabilizes a mesenchymal-like state in these cells.

The context-dependent duality of TAM function in *Drosophila* with plasmatocytes exhibiting pro-tumorigenic functions in some genetic contexts and anti-tumorigenic functions in others (Hirooka et al., 2025; Parisi et al., 2014; Voutyraki et al., 2023) mirrors the phenotypic plasticity of mammalian TAMs and underscores the importance of the local tumor microenvironment in shaping immune cell behavior (Murray et al., 2014; Pittet et al., 2022). Recent single-cell sequencing of *Drosophila* TAMs in tumor contexts has revealed transcriptional heterogeneity analogous to the functional spectrum observed in mammalian TAM populations (Khalili et al., 2023; Yarikipati & Bergmann, 2026), raising the possibility that distinct TAM subpopulations at the leading edge versus within the tumor mass play non-redundant roles in supporting metastatic colonization. Our spatial quantification showing TAM distribution across both involved and un-involved tumor regions is consistent with this model, and future single-cell or best spatial transcriptomic analysis of TAMs in this system may reveal functionally specialized subsets.

### *Drosophila* for modelling tumor cell states in organ metastasis

A striking feature of *lgl*^-/-^ tumor metastasis to the adult brain is the dynamic and compartment-specific regulation of cadherin expression. Metastasized tumor cells exhibited robust DE-cadherin expression in both punctate/cortical and dense plaque-like accumulations at cell-cell interfaces consistent with adherens junctions a pattern not observed in control NSCs, where DE-cad localizes without forming canonical junctions (Almeida & Bray, 2005; Banach-Latapy et al., 2023). Even more striking is the widespread upregulation of DN-cadherin in tumor cells, despite their NSC origin: NSCs and their progeny are normally DN-cad-negative, with DN-cad restricted to mature neurons where it mediates synaptic adhesion (Iwai et al., 2002; Kurusu et al., 2012).

The co-expression of DE-cad and DN-cad in distinct subcellular compartments including dense plaque-like accumulations confirmed by electron microscopy as electron-dense junctional structures is directly analogous to the cadherin switch that defines epithelial-to-mesenchymal transition in mammalian cancers, where downregulation of E-cadherin and upregulation of N-cadherin facilitates invasive and migratory behavior (Mrozik et al., 2018; Theveneau & Mayor, 2012). In glioblastoma specifically, aberrant N-cadherin expression drives invasion along white matter tracts and perivascular spaces, and N-cadherin-mediated heterotypic junctions between tumor cells and cancer-associated fibroblasts are important facilitators of collective invasion in carcinomas (Mrozik et al., 2018). Whether DN-cad upregulation in *lgl*^-/-^ tumor cells at the brain surface reflects a bona fide EMT-like transcriptional reprogramming, a lineage de-differentiation toward a neuronal identity, or a functional adaptation to the brain microenvironment remains an important open question. Given that *Drosophila* offers both the ability to genetically manipulate cadherin expression with cell-type specificity and the imaging access to track tumor cell behavior in real time, this system is uniquely positioned to dissect the causal relationship between cadherin switching and invasive capacity at metastatic sites.

The spatial cell morphology data reported here systematic differences in cell aspect ratio between un-involved, orthogonally invasive, and laterally spreading tumor cell populations provides strong evidence for a leader-follower model of collective invasion at the brain surface. In this framework, elongated leader cells at the invasive front, characterized by mesenchymal-like morphology and higher aspect ratios, guide the directional movement of rounder follower cells that contribute mechanical force through passive pushing (Friedl & Mayor, 2017; Yamamoto et al., 2023). The co-expression of both DE-cad and DN-cad with adherens junction-like contacts between adjacent cells is consistent with the requirement for maintained cell-cell adhesion that is a hallmark of collective, as opposed to single-cell, invasion modes.

Together, our findings establish the adult *Drosophila* brain as a tractable *in vivo* platform for dissecting the cellular dynamics of tumor-barrier and tumor-immune interactions during metastatic colonization at single-cell resolution. Future studies leveraging this platform to identify the molecular signals driving TAM recruitment, the function of BM, and the molecular basis of leader cell specification at metastatic sites have the potential to reveal conserved mechanisms governing organ metastasis.

## MATERIAL AND METHODS

### *Drosophila* genetics and fly husbandry

Flies were raised on the standard Bloomington stock center food recipe (LabExpress, Ann Arbor, MI, USA). Flies were cultured in an incubator maintained at 25°C and 40±5% relative humidity. No distinction was made between male and female larvae in the experiments conducted with larval tissue. For Gal80ts experiments, flies were raised at 16°C until adulthood and then transferred to 29°C to activate Gal4.

#### Drosophila strains

*lgl^4^-Frt40*, *Frt40* MARCM, *Vkg-GFP* (Bl. 98343), *Trol-GFP*, ZCL1700) (Morin et al., 2001), PNG-Gal4 (Bl. 40436), SPG-Gal4 (Bl. 50472), Cortex-Gal4 (Bl. 45784), 10XUAS-IVS-mCD8::GFP (Bl. 32185) and (Bl. 32186), UAS-10X-IVS-mCD8-RFP (Bl. 32219), UAS-*rpr* (Bl. 5824), UAS-*rpr, hid* (Gift from Tin Tin Su, University of Colorado Boulder).

### Allograft transplantation protocol

Transplantation was carried out by adapting the protocol described in (Rossi & Gonzalez, 2015). 3- to 5-day-old *w^11-18^* virgin female flies were used as the host flies for primary transplantation and obtaining the T0 tumor. Following primary transplantation, the tumor mass was further amplified by serially transplanting T0 to obtain T3-T5 stage tumors, refer to in our previous paper (Khan & Rusan, 2025). We then utilized the T3-T5 tumor for injecting into 3- to 5-day-old virgin host female flies of a desired genotype. Following transplantation the flies were maintained at 25°C incubator.

### Immuno-histochemistry

A tissue-specific fixation and immunostaining protocol was followed. Dissected tissues were fixed in 4% formaldehyde and then incubated in blocking buffer containing 0.5% bovine serum albumin (BSA) in 0.1% PBST (1X PBS with 0.1% Triton X-100). Primary antibody incubation was carried out overnight at 4°C, followed by three washes in 0.1% PBST at room temperature. The following primary antibodies were used: mouse anti-NimC1/2 (1:50; gift from István Andó at the Institute of Genetics, HUN-REN Biological Research Centre, Szeged, Hungary), and rat anti-Elav (9F8A9, DSHB, 1:10). After washing, tissues were incubated with Alexa Fluor 488-, 564- or 647-conjugated secondary antibodies (1:250) at room temperature, followed by three additional washes in 0.1% PBST. This was followed by incubation with an HSC NuclearMask (Invitrogen H10325, 1:2000) for 10 minutes, followed by a final wash with 0.1% PBST. If stained for F-actin, tissues were incubated with Phalloidin-Atto-647 (Sigma 65906, 1:1000) for 15 minutes, then washed with 0.1% PBST. Samples were then rinsed in 1X PBS and mounted on slides using Aqua-Poly/Mount (Polysciences 18606-5) and cured overnight at room temperature. A 35-40 µm heigh nail polish spacers was used to maintain a similar level of comparison of brains across samples.

### BBB permeability assay

To assess blood-brain barrier (BBB) integrity, a fluorescent dextran permeability assay was performed as previously described with minor modifications (Love & Dauwalder, 2019). Flies were anesthetized and injected in the thorax with ∼100 nL of 25 mg/mL 10 kDa lysine-fixable Dextran, Texas Red™ (Thermo Fisher Scientific, D1863). Following injection, flies were maintained at 25°C overnight to allow systemic circulation of the tracer dye. For SPG ablation controls, *SPG>rpr* flies were shifted to 29°C for 5 days prior to injection to induce cell death of SPG cells. Control flies (*w^1118^* injected with media) and tumor-bearing flies (*lgl^-/-^*; hosts in *w^1118^* background) were analyzed at Day 6 post-transplantation. After overnight incubation, heads were then removed and fixed for 30 minutes in 4% formaldehyde. Brains were dissected in 4% formaldehyde, fixed for additional 30 minutes, washed in PBS, and mounted in Aqua-Poly/Mount medium on glass slides with 35-40 µm spacers.

Samples were imaged using a Zeiss LSM 980 laser-scanning confocal microscope with a 20X objective (0.8 NA). All three samples were imaged using identical laser and gain settings. Intensity measurements were performed by averaging mean intensity of multiple ROIs from the central brain and optic lobes (Figure S5). Bright signal in the center of the brain was omitted from intensity measurements.

### Imaging and image processing

Image acquisition was performed using Zeiss laser-scanning confocal microscopes (LSM 880 and 980). For imaging entire adult brains, samples were acquired with a 60X objective (1.4 NA) using tiling settings with 10% overlap between adjacent tiles and a Z-step size of 1 µm for full brain. Image stitching was performed post-acquisition using ZEN Blue software. For high-resolution imaging of specific tumor or brain regions, regions of interest (ROIs) were selected following acquisition of the whole-brain tiled images.

Raw image data were processed using Fiji, and appropriate Z-sections were subjected to maximum-intensity projection. Adobe Creative Cloud software (Photoshop and Illustrator) was used to adjust image intensity and prepare figures for publication.

### FIB-SEM sample preparation and imaging

Prior to imaging, samples were fixed in 2.5% glutaraldehyde, post-fixed in osmium tetroxide, dehydrated through a graded ethanol series, and embedded in resin. Samples were mounted on aluminum SEM stubs using conductive silver paint and sputter-coated with a ∼10 nm layer of gold to minimize charging.

Samples were imaged using a dual-beam FIB-SEM system Zeiss Crossbeam 540 equipped with a gallium (Ga⁺) ion source. The region of interest (ROI) was identified by SEM imaging, and a protective platinum (Pt) pad approx. 20 × 40 µm was deposited over the ROI via ion-beam-assisted deposition to preserve the surface during milling. A coarse trench was milled adjacent to the ROI using a high ion-beam current 5000pA at 30 kV to expose the cross-sectional face, followed by fine polishing at a reduced current 1-3 nA. For volume acquisition, serial sectioning was performed by alternating ion-beam milling and SEM imaging (slice-and-view). Drift correction and focus/stigmation adjustments were applied automatically throughout acquisition. The resulting image stacks were aligned, registered, and processed using Fiji/ImageJ.

### Tumor Volume measurements

Full brain stitched images were exported using BIO-FORMAT to OME.TIFF from CZI. OME to preserve meta data information. OME.TIFF files were imported to Imaris 10.2.0 version software. Using surpass volume rendering module 3D volumes of a given channel was rendered and the volumetric measurements in µm^3^ were obtained from the statistics tab.

### Brain deformation measurements

Dissected adult brains were mounted on glass slides with approximately 35-40μm high nail polish spacers. This resulted in an even flattening of the brains across samples. The samples were imaged on a Zeiss-980 laser scanning microscope using a 63X objective, 1X digital zoom, and a 1µm Z-step size. Images were tiled with a 10% overlap and stitched post-acquisition using Zen Blue software.

#### Quantification of the extent of tumor invasion

Brains were optically sliced with a ∼5 µm step size. The middle frame was used as the reference and top −5 µm and bottom +5 µm were used as experimental frames (Figure 1C and S2). *w* represents a point on the brain outline in the experimental frame, and Δ(*w*) denotes the short distance from *w* to the reference outline, used as a measure of deformation magnitude (Figure S2, orange arrow). To compensate for minor variability in brain sizes across experiments, we computed Δ(*w*) relative to the reference brain circumference. The mean deformation Δ was computed by taking the average of Δ(*w*) evaluated at N=1000 points *w* uniformly distributed over the circumference of the experimental frame.

### Statistical analysis

Statistical analysis was performed using GraphPad Prism 10. All graphs display the mean ± SEM with individual data points shown. Statistical significance between two groups was assessed using a two-tailed Student’s t-test. For comparisons across multiple time points or genotypes, one-way ANOVA with Tukey’s or Dunnett’s multiple comparisons test was used. P < 0.05 was considered statistically significant. Significance levels are indicated as follows: ****P < 0.0001, ***P < 0.001, **P < 0.01, and *P < 0.05.

## ACKNOWLEDGEMENTS

We thank Xufeng Wu, and Christian Combs from the NHLBI Light Microscopy Core, Valentina Baena Echeverri and Zulfeqhar Syed from the NHLBI electron microscopy core, Nikhil Karthik (American Physical Society) for help with writing algorithm for quantifying brain deformation, members of Rusan lab, Alexandar Kelly lab (National Cancer Institute), and Takashi Akera lab (NHLBI) for helpful discussions, the Bloomington *Drosophila* Stock Center for fly stocks, and the Developmental Studies Hybridoma Bank for antibodies. This work was supported by the Division of Intramural Research at the National Heart, Lung, and Blood Institute (ZIAHL006126 to NMR).

## DISCLAIMER

This research was supported by the Intramural Research Program of the National Institutes of Health (NIH). The contributions of the NIH author(s) are considered Works of the United States Government. The findings and conclusions presented in this paper are those of the author(s) and do not necessarily reflect the views of the NIH or the U.S. Department of Health and Human Services.

## DATA AVAILABILITY STATEMENT

All relevant data can be found within the article and its supplementary information. The DOI for all raw data is 10.25444/nhlbi.32807480 (Data provided following final acceptance of manuscript).

---

**Figure S1.**
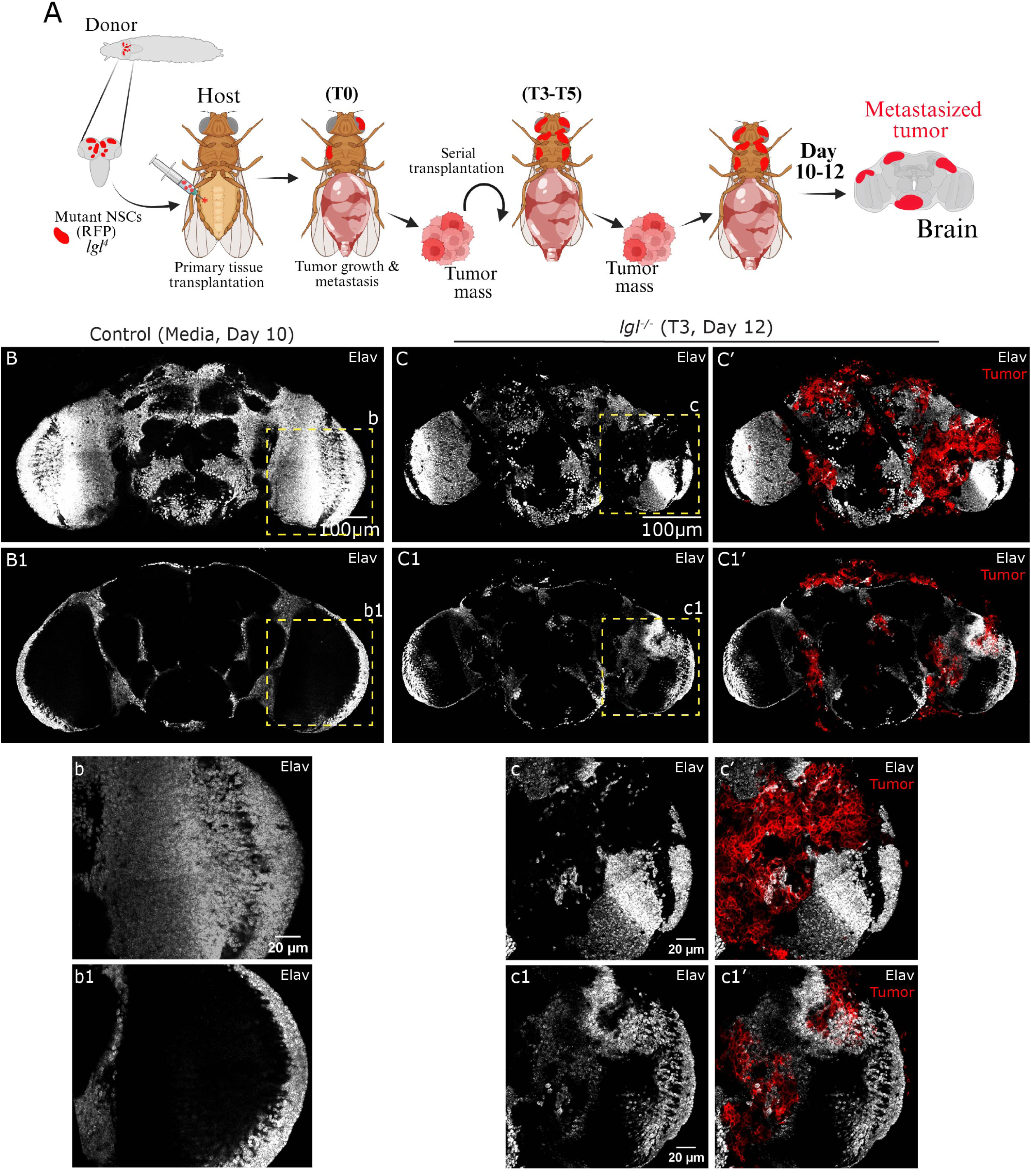
*lgl*^-/-^ tumors deform and disrupt neuronal tissue organization in the *Drosophila* brain. **(A)** Schematic illustration of the tumor transplantation model. *lgl^-/-^*tumor cells are dissected and injected into the abdomen of host *Drosophila*. Tumors grow progressively within the host and ultimately metastasize to the brain. **(B-B1)** Representative confocal z-stack images of a control (Media, Day 10) brain at two z-levels showing Elav-positive neurons (grey scale). **(b-b1)** High-magnification images of the boxed regions in **B** and **B1**, respectively, showing Elav distribution in control brains. **(C-C’)** Representative confocal z-stack images of an *lgl^-/-^* brain (T3, Day 12) at the top z-level showing Elav (grey) alone (**C**) and merged with tumor cells (red) (**C’**). **(C1-C1’)** Mid z-level images of the same *lgl^-/-^* brain showing Elav alone (**C1**) and merged with tumor (red) (**C1’**). **(c-c’)** High-magnification images of the boxed region in **C** showing Elav (grey) alone and merged with tumor cells (red), respectively, illustrating tumor disrupting the neuronal cell cortex. **(c’-c1’)** High-magnification images of the boxed region in **C1** showing Elav (grey) alone and merged with tumor (red) at a deeper z-section.

**Figure S2.**
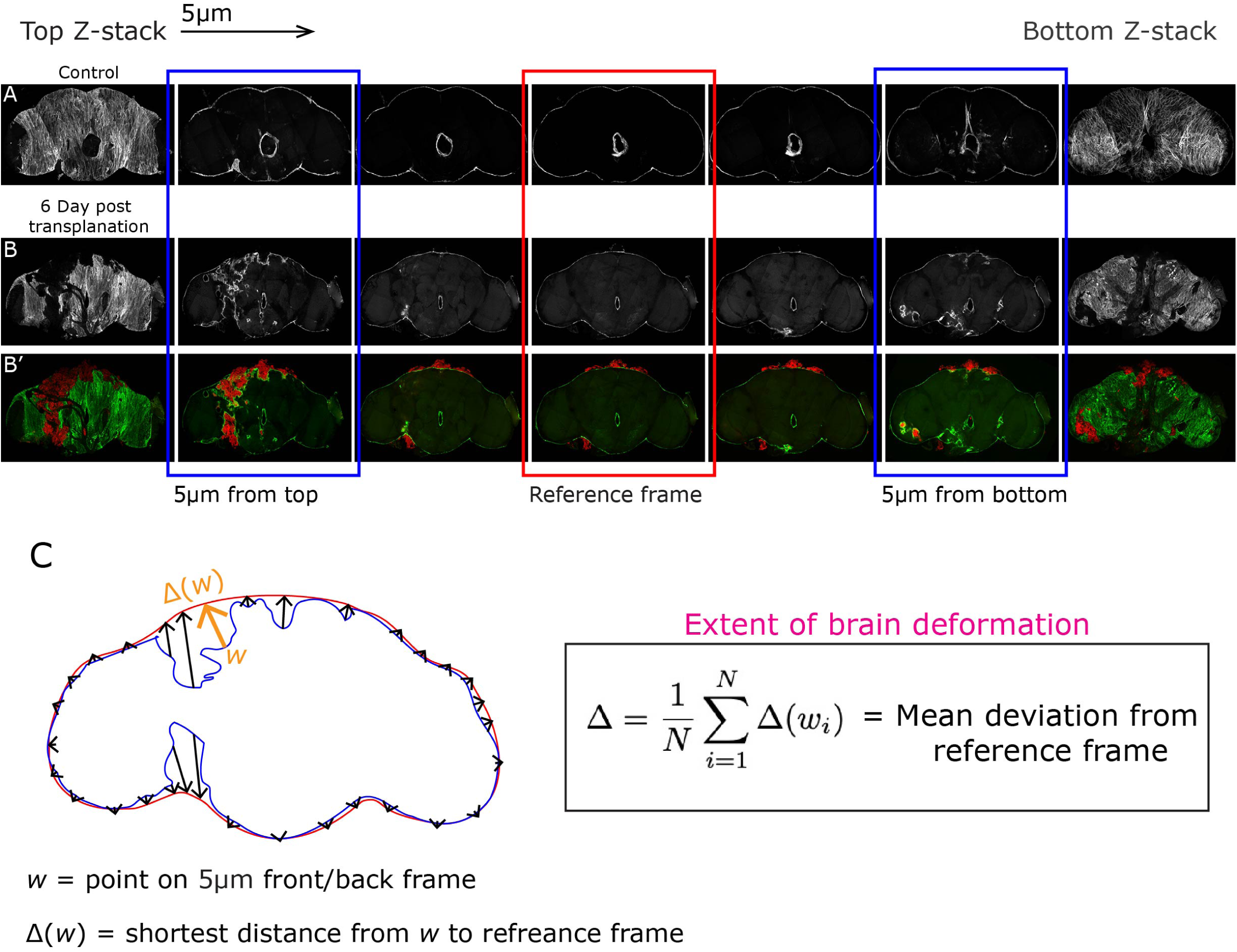
Schematic of the brain deformation quantification pipeline using serial confocal z-stack. **(A)** Representative serial confocal z-stack images spanning from the top to the bottom z-plane of a *Drosophila* brain, showing brain outline marker PNG-Gal4 driving mCD8-GFP. The central image highlighted by the red box denotes the reference frame (middle z-section) used as the fixed anatomical reference point. Blue boxes indicate the experimental frames at the top and bottom z-sections used for deformation comparison. **(B-B’)** Corresponding serial z-stack images of an *lgl^-/-^* tumor-bearing brain shown in **B**, and a merged view of PNG glia (green) and tumor cells (red) in **B’**, across the same z-planes. The red-boxed central image represents the reference frame. Blue boxes highlight the top and bottom experimental frames. **(C)** Schematic diagram illustrating the method used to quantify the extent of brain deformation. The brain outline from the reference frame (middle z-section, red) is superimposed onto the outline from an experimental frame (blue). *w* represent point on brain outline on experimental frame, and Δ(*w*) denotes the deviation of the experimental frame outline from the reference, used as a measure of deformation magnitude (see methods).

**Figure S3.**
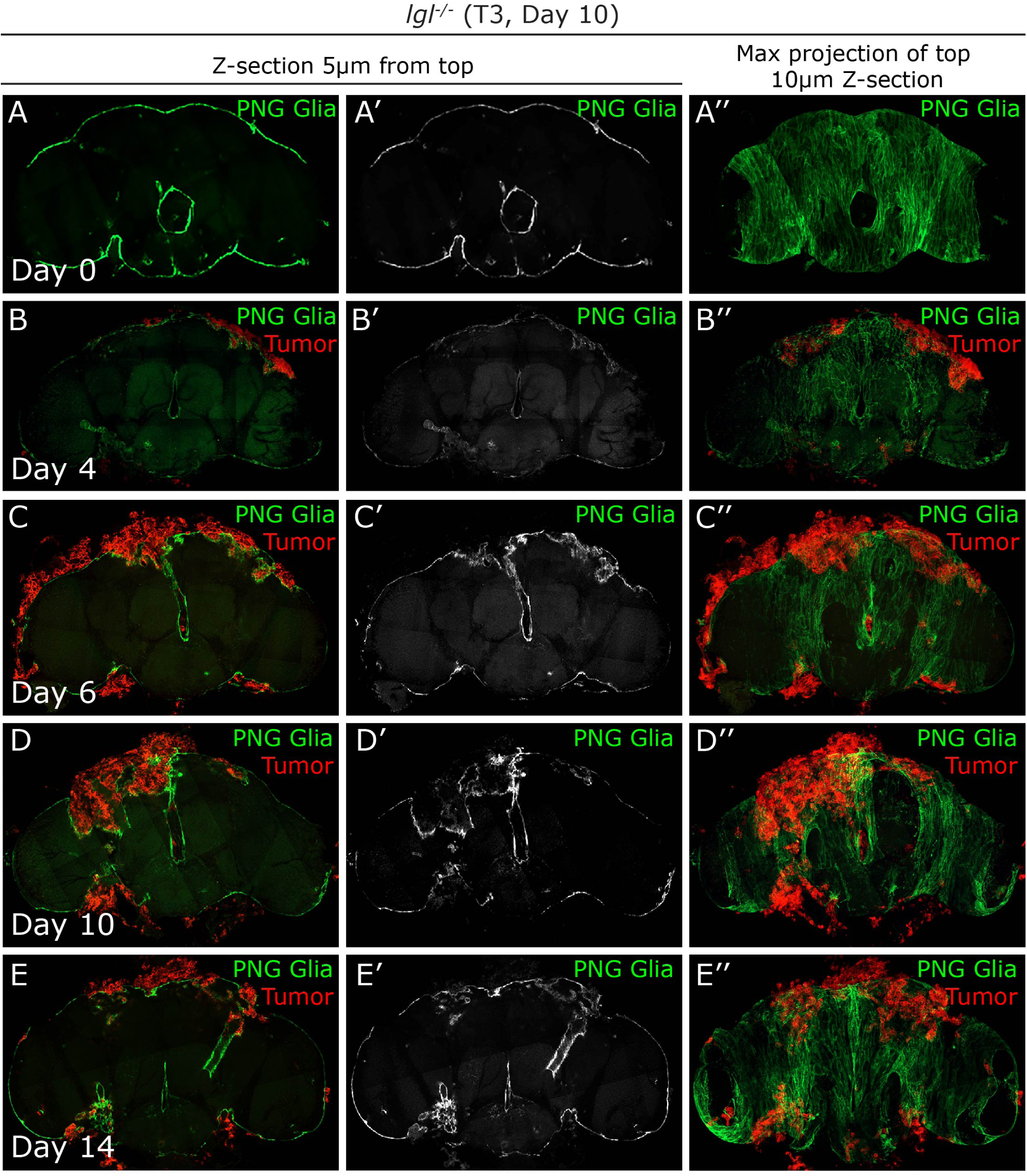
Progressive disruption of brain surface by *lgl*^-/-^ tumors over days. Representative confocal images brain showing PNG Glia (green) −5 µm z-section from the top, and max projection of top 10 µm z-sections at Day 0 **(A-A’’)**, Day 4 **(B-B’’)**, Day 6 **(C-C”)**, Day 10 **(D-D”)** and Day 14 **(E-E”)** after T3-*lgl^-/-^* tumor (red) injection.

**Figure S4.**
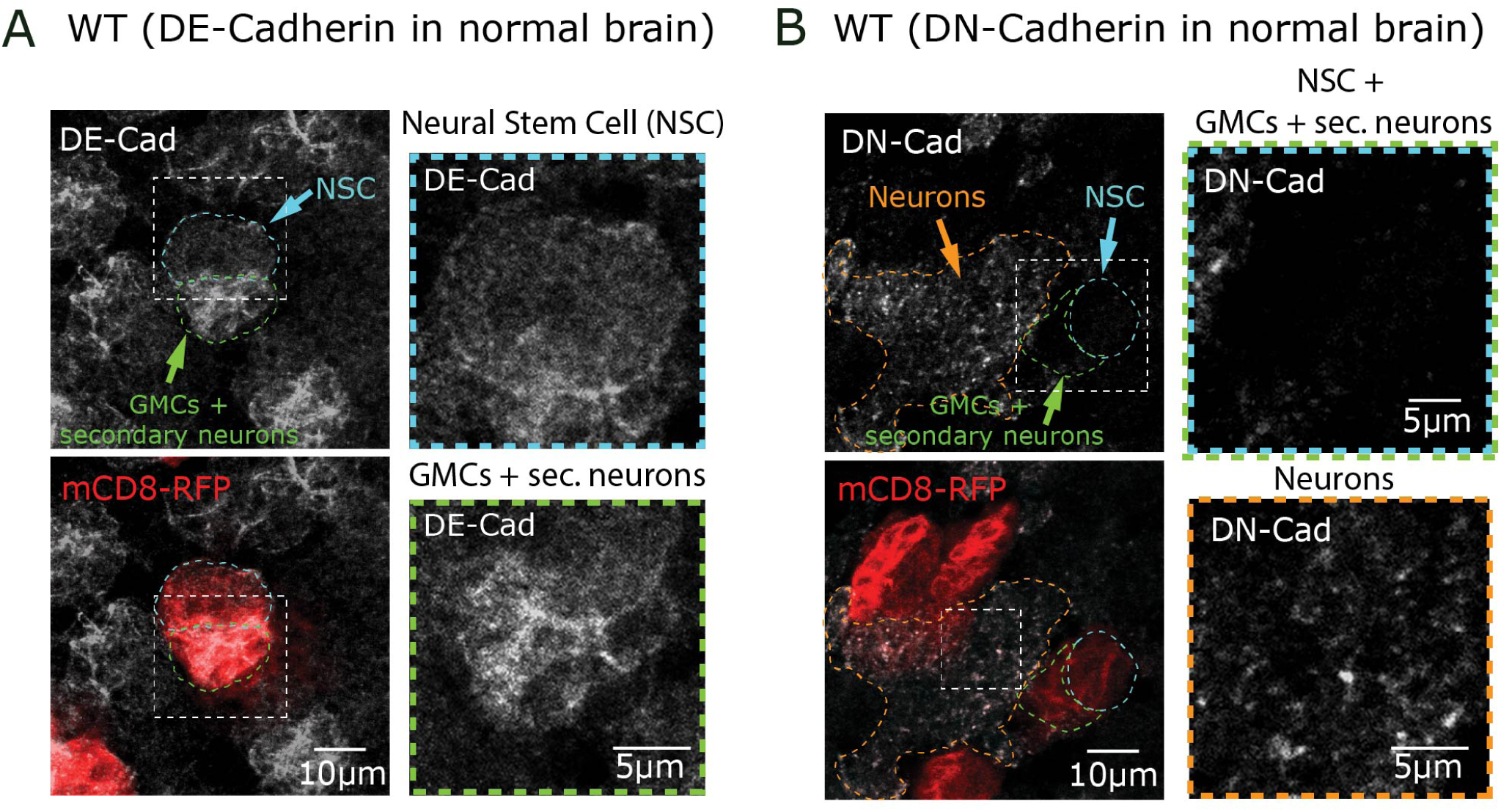
DE-Cadherin and DN-Cadherin localization in wild-type brains cells. **(A)** Representative confocal images of a wild-type (WT) brain stained for DE-Cad (grey scale) with MARCM clone (mCD8-RFP, red) labeling neural stem cell lineage. Image showing DE-Cad distribution with a cyan outline indicating a neural stem cell (NSC, cyan arrow) and a green outline indicating ganglion mother cells (GMCs) and secondary neurons (GMCs + sec. neurons, green arrow). High-magnification insets showing DE-Cad localization in the NSC (cyan box) and GMC + secondary neuron region (green box). **(B)** Representative confocal images of a WT brain stained for DN-Cad (grey). Image showing DN-Cad distribution with an orange outline indicating neurons (orange arrow), a cyan + green outline indicating an NSC + GMCs (cyan + green arrow). High-magnification insets showing DN-Cad localization in the NSC + GMCs (cyan +green box) and secondary neuron region (orange box).

**Figure S5.**
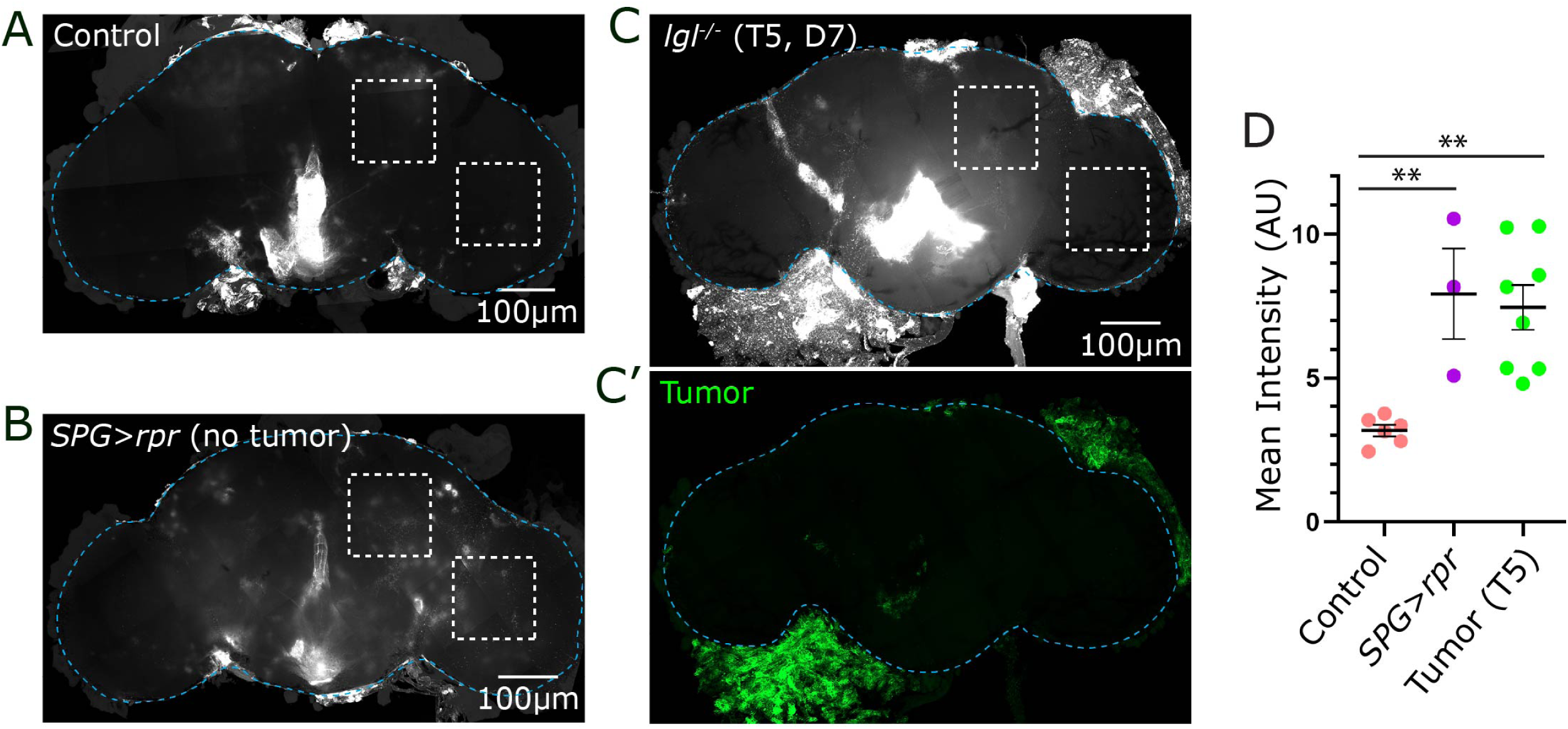
*lgl^-/-^* tumor metastasis compromises functional integrity of the BBB. **(A)** Representative confocal image of a control brain showing dextran signal. **(B)** Representative confocal image showing dextran signal of a *SPG>rpr* brain (no tumor), in which subperineurial glia are genetically ablated via expression of the pro-apoptotic gene *rpr* under the *SPG*-Gal4 driver. **(C-C’)** Representative confocal images of an *lgl^-/-^*tumor-bearing brain (T5, Day 7), **C** shows dextran signal and **C’** shows the corresponding tumor channel (green). White dashed boxes in (A-C) indicate ROIs used for mean intensity quantification. **(D)** Quantification of mean fluorescence intensity (arbitrary units, AU) measured within the ROIs shown in (A-C) for control, *SPG>rpr* no-tumor, and *lgl^-/-^* tumor T5. Data are presented as mean ± SEM, one-way ANOVA with Tukey’s multiple comparisons test was used. P < 0.05 was considered statistically significant. Significance levels are indicated as follows: ****P < 0.0001, ***P < 0.001, **P < 0.01, and *P < 0.05.

**Figure S6.**
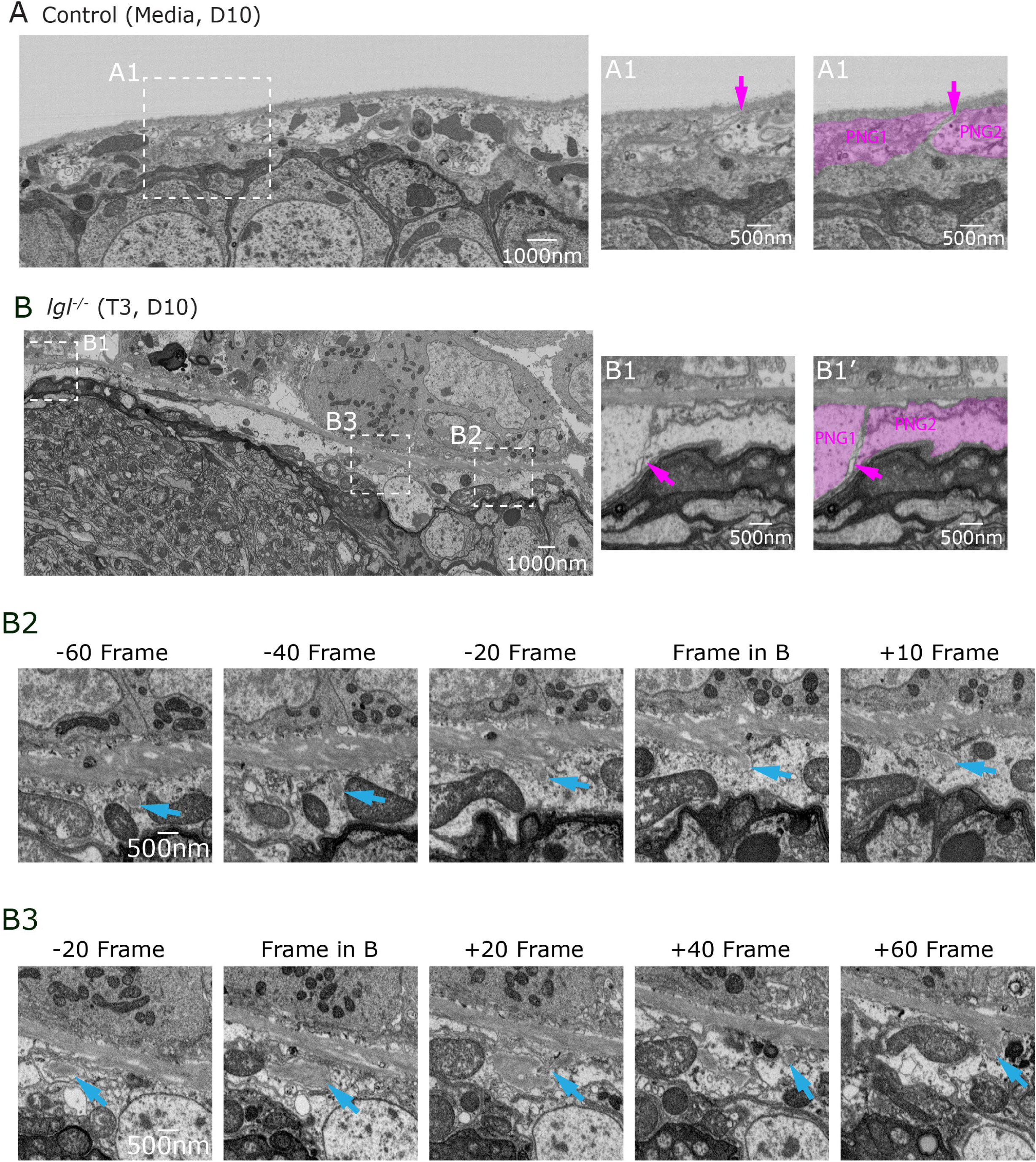
Ultrastructural analysis of the BBB at metastatic site. **(A)** FIB-SEM overview image of a control (Media, Day 10) brain showing the intact BBB in cross-section. **(A1-A1’)** High-magnification FIB-SEM image of the boxed region in A showing the normal ultrastructural organization of the PNG cells, the magenta arrow indicates intact PNG-PNG contact. **(B)** FIB-SEM overview image of an *lgl^-/-^* brain (Transplant 3, Day 10). **(B1-B1’)** High-magnification FIB-SEM image of the boxed region in B the magenta arrow indicates loss of PNG-PNG contact. **(B2)** Serial FIB-SEM sections through the region boxed as B2 in B, shown at frames −60, −40, −20, the reference frame (Frame in B), and +10, spanning, blue arrows through consecutive sections marks vesicle containing ECM material in continuity with BM. **(B3)** Serial FIB-SEM section series through the same region at frames −20, the reference frame (Frame in B), +20, +40, and +60, blue arrows through consecutive sections marks vesicle containing ECM material in continuity with BM.

**Figure S7.**
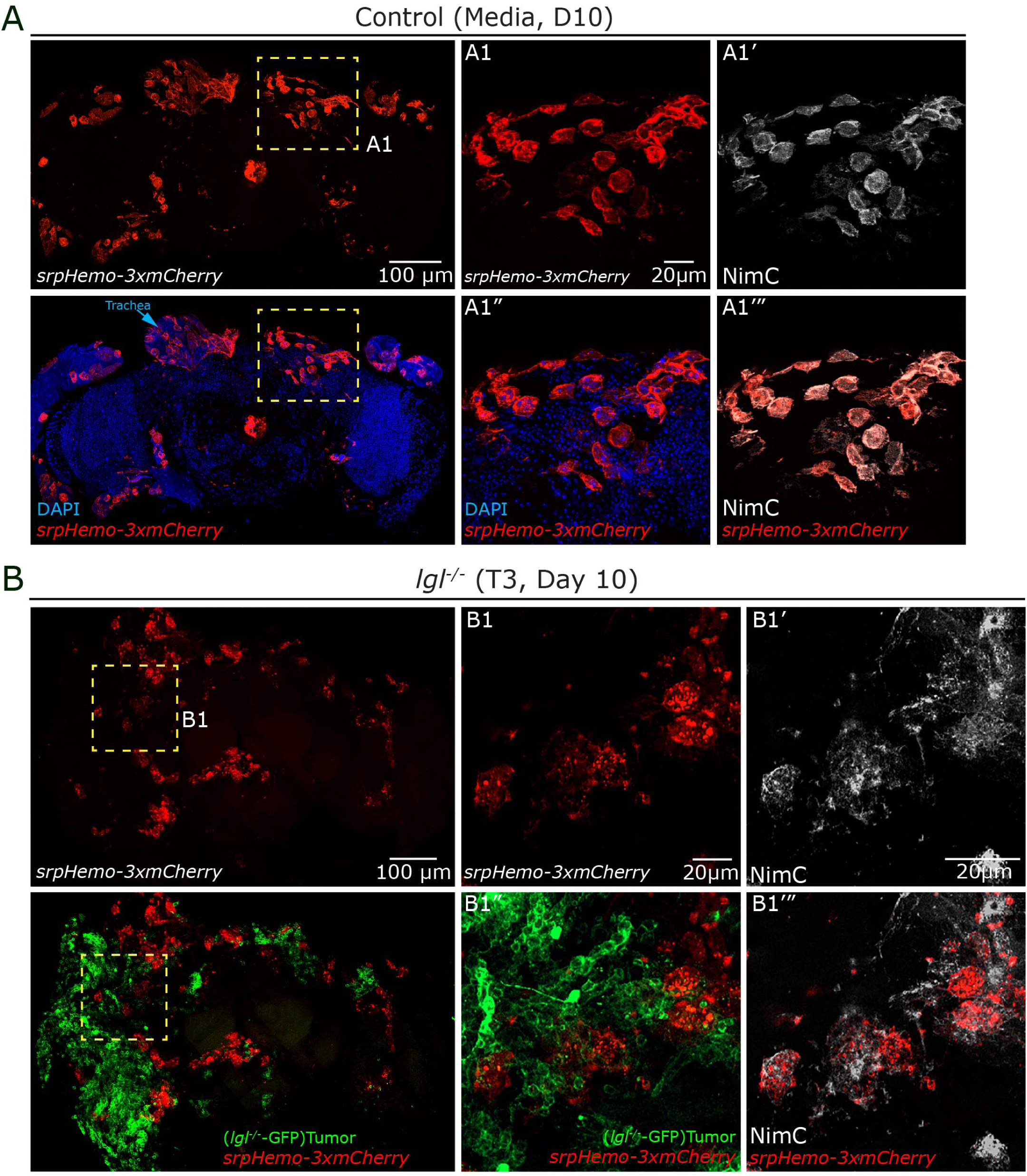
*lgl^-/-^* tumors recruit activated macrophages. **(A)** Representative images of a control (Media, Day 10) brain showing hemocytes labeled with *srpHemo-3xmCherry* (red). **(A1)** High-magnification image of the boxed region showing *srpHemo-3xmCherry*-positive hemocytes (red). **(A1’)** Corresponding NimC channel (grey scale) confirming hemocyte identity in the same region. **(A1”)** High-magnification merged image showing DAPI (blue) and *srpHemo-3xmCherry* (red) hemocytes. **(A1”’)** Merged image of NimC (white) and *srpHemo-3xmCherry* (red) confirming co-labeling of hemocytes by both markers in control brains. **(B)** Representative confocal images of an *lgl^-/-^* brain (T3, Day 10). **(B1)** High-magnification image showing *srpHemo-3xmCherry*-positive hemocytes (red) in the tumor-proximal region. **(B1’)** Corresponding NimC channel (grey scale) showing hemocyte distribution in the same region, with enlarged and irregularly shaped cells consistent with an activated plasmatocyte morphology. **(B1”)** High-magnification merged image of *lgl^-/-^* -GFP tumor cells (green) and *srpHemo-3xmCherry* hemocytes (red) showing hemocytes closely associated with tumor cells. **(B1’’’)** Merged image of NimC (white) and *srpHemo-3xmCherry* (red) confirming co-labeling of hemocytes by both markers in tumor-bearing brains.

**Figure S8.**
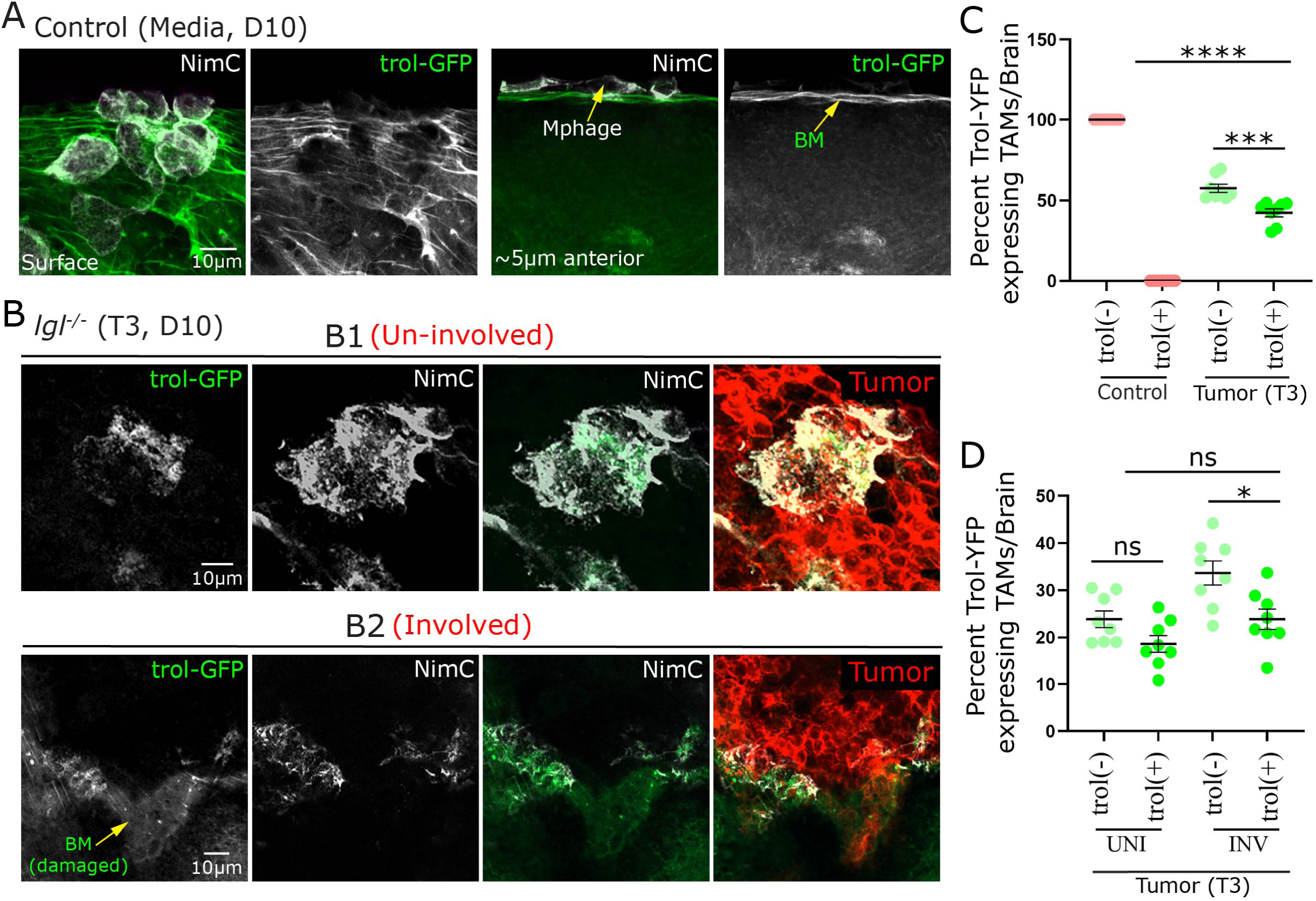
TAMs positive Perlecan/Trol are uniformly distributed in tumor-involved regions of *lgl*-deficient brains. **(A)** Representative confocal images of a control (Media, Day 10) brain showing hemocyte-associated Trol expression. Left two panels, surface view showing NimC-positive hemocytes (grey scale) and *trol-GFP* signal (green) at the brain surface. Right panels, Z-section approximately 5 µm anterior to the surface showing a NimC-positive hemocyte (Mphage, yellow arrow) and *trol-GFP* signal (green) localized along the basement membrane (BM, yellow arrow). In control brains, *trol-GFP* expression is restricted to the BM with minimal hemocyte-associated signal. **(B)** Representative confocal images of an *lgl^-/-^* brain (Transplant 3, Day 10) showing Trol expression in TAMs across two distinct regions. **(B1)** (Un-involved region) High-magnification images showing *trol-GFP* (grayscale, left), NimC (grayscale (white, second panel), NimC in green (green, third panel), and tumor cells (red, right) in a region without direct tumor contact. **(B2)** (Involved region), High-magnification images showing *trol-GFP* (grayscale, left), NimC (grayscale (white, second panel), NimC in green (green, third panel), and tumor cells (red, right) in a region with direct tumor contact. **(C)** Quantification of the percentage of *trol-GFP*-expressing tumor-associated macrophages (TAMs) per brain in control and *lgl^-/-^* T3 transplanted, subdivided by *trol* expression status [trol(−) and trol(+)]. **(D)** Quantification of the percentage of *trol-GFP*-expressing TAMs per brain in un-involved (UNI) and tumor-involved (INV) regions of Tumor T3 brains, subdivided by trol expression status [trol(−) and trol(+)]. Data are presented as mean ± SEM, one-way ANOVA with Tukey’s multiple comparisons test was used. P < 0.05 was considered statistically significant. Significance levels are indicated as follows: ****P < 0.0001, ***P < 0.001, **P < 0.01, and *P < 0.05.

## Notes

### Competing Interest Statement

The authors have declared no competing interest.

